# An open source microfluidic sorter for *Caenorhabditis* nematodes

**DOI:** 10.1101/780502

**Authors:** Nadine Timmermeyer, Stephen A. Banse, Joseph H. North, Matthew Sottile, Shawn R. Lockery, Patrick C. Phillips

**Author notes:** These authors contributed equally to this work (Nadine Timmermeyer, Stephen A. Banse).

## Abstract

Rapid and accurate sorting of biological samples is extremely useful in a wide variety of applications. The model nematode *Caenorhabditis elegans* lends itself to automated fluid-flow based sorting because of its ability to live in aqueous solutions. Here, we build upon previous developments to construct a microfluidic device capable of sorting individuals based on a variety of characteristics, with a specific application toward differentiating fluorescently marked individuals. We find that our new design generates highly repeatable pools of sorted individuals. In general, there tends to be a tradeoff between precision and speed that can be optimized based on several different factors. Importantly, sorting does not decrease offspring production, and individuals can be sorted multiple times for increased precision. We provide detailed parts lists, schematics and software to allow implementation of these methods in other laboratories. Our results demonstrate that custom-made sorter chips can be a flexible and versatile addition to the nematode experimental toolbox.

## Introduction

*Caenorhabditis elegans* is one of the most popular animal model systems in biological research [1]. While its small size enables maintenance of large populations, it can be a challenge when those large populations need to be subdivided by visual phenotype. In standard practice, individuals that need to be sorted according to visual phenotype have to be identified by the experimenter and transferred by hand. The time-consuming nature of manual transfers limits the number of nematodes that can be processed at a given time, therefore limiting the population size, replicate number and/or treatments for any given project. This limitation can reduce both the scale and feasibility of certain classes of experiments (e.g., saturating mutagenesis, experimental evolution).

One possible solution to the limitations imposed by manual animal transfer is to automate the sorting process using fluid-flow based nematode sorters. The basic premise of any nematode sorter is to capitalize on the ability of *Caenorhabditis* nematodes to survive in an aqueous environment to accomplish the four phases in sorting: (1) hold the population to be sorted in a reservoir where they are primed to move through the device, (2) move single animals into the optical path of an imaging system, (3) capture visual information about the individual and make a determination of phenotype, (4) redirect the liquid flow to move the individual into the desired pool, and then repeat. For a number of years, *C. elegans* researchers have been able to use a commercial product, the Union Biometrica COPAS, for this purpose. The COPAS FP (and now BioSorter and COPAS Vision) is a large particle flow cytometer capable of sorting small organisms like *C. elegans*. A major advantage of the COPAS is the extremely large number of samples that can be processed per hour via the use of a laser detection system. However, the large purchase price and maintenance costs of this instrument limits access for most labs. Additionally, the binary nature of sorting, selected or discarded, in the COPAS system limits the complexity of sorting, and requires multiple sorts of a population if more than one subpopulation is to be selected. Further, the laser-based system and constant motion of the worms precludes some forms of sample characterization that are possible with conventional optical microscopy.

An alternative to purchasing a commercial large particle flow cytometer is to use soft lithography to generate custom-fabricated silicon chips that automate worm handling while paired with the imaging hardware already present in most *C. elegans* labs. Soft lithography fabricated polydimethylsiloxane (PDMS) chips have become very popular in various applications in physics and chemistry and are playing an increasingly important role in biology [2]. PDMS is non-toxic, oxygen permeable, and optically transparent, making it particularly attractive for use with nematodes. PDMS-based microfluidic chips have been used in a wide range of nematode research applications [3,4], including image-based screening [5–10], assaying lifespan [11], stress resistance [12], and reproduction [13], analyzing neurophysiology and behavior [11,14–19], and sorting animals based on size and/or behavior [20]. In addition to these applications, nematodes can be sorted in microfluidic devices based on visual markers [5,21].

Microfluidic sorters that sort based on optical information are typically constructed of at least two layers, a flow layer and a control layer. The flow layer contains the worm inlet, multiple worm outlets, and channels connecting the inlets to the outlets. The control layer enables directed movement of worms through the flow layer channels using control channels that physically overlap the flow channels while being separated by a thin, flexible PDMS septum. When control channels are pressurized the septum acts as a valve by stretching into the overlaid flow channel, blocking the flow channel at any point of intersection. This two-layer approach has been used in two major design types for phenotypic sorting of nematodes, suction-capture and channel capture designs. The suction-capture designs use a circular channel path in which individuals flow through the device in a cyclical pattern and are periodically captured by a suction device [22]. Excess individuals flow out of the system while the captured animal is processed. This design requires pressurized air to drive the flow of individuals, as well as a vacuum source to suction-capture individual nematodes. In channel-capture designs, individuals flow down a narrow channel in which they become trapped for the duration of image acquisition and analysis, and subsequently are flushed into an outlet channel of choice [6]. These designs rely solely on pressurized air for overall fluid control. This approach can be further augmented by adding a third layer of channels (with additional complexity) for a cooling fluid to immobilize the nematode, facilitating higher magnification or longer exposure time imaging [21].

Here, we build upon previous channel-capture devices to develop a microfluidic nematode sorter that is both robust and easy to use. In particular, we modified a previously published channel-capture design [5] by adding a loading valve to prevent two individuals from simultaneously entering the imaging channel, a flush channel to move individuals through the system with more precision, and an additional outlet channel to increase sorting options. The additional outlet channel enables the usage of a default sorting outcome of “no choice”. Additionally, we have developed custom MATLAB (MathWorks, Inc.) software which sorts individual nematodes based on either experimenter determination (i.e., scored visually and directed to the designated channel via keyboard control) or by automated image acquisition and analysis of fluorescent markers. The software implementation is highly flexible, being independent of the type of camera, microscope, and imaging software used. Importantly, all designs and details related to the production of these devices are provided here.

We demonstrate the utility of our approach by applying it to three sorting scenarios using *C. elegans*. In the first scenario (Fig. 1, Scenario A), we sorted developmentally equivalent sub-populations of nematodes that expressed either of two distinct, bright, fluorescent markers. In the second scenario (Fig. 1, Scenario B) we sorted developmentally different subpopulations (males and hermaphrodites) using sex-specific expression of GFP markers. This application represented increased sorting difficulty in all four phases of sorting by introducing developmentally coupled reporter expression, as well as potential body-size and behavioral differences. Many applications including experimental evolution and sex specific tissue collection require that large number of males and hermaphrodites (or females) be separated from each other, preferably before they become sexually mature. For the third scenario (Fig. 1, Scenario C), we mimic the conditions for separating transgenic from non-transgenic individuals by sorting GFP marked from non-marked nematodes. This scenario represents a sorting challenge in the third phase of sorting because one phenotypic class is invisible to the imaging system. Since this latter approach might be applied in situations where the desired individuals exhibited either the gain or loss of marker expression, we included different ratios of the two marker classes and analyzed the relationship between starting ratio and resulting precision. Across all three Scenarios, we find sorting precision to be high, but not perfect. Secondary sorting is an option and does help to increase precision. With the increasing ease and availability of designing and manufacturing microfluidic chips within individual labs, our new device should prove to be a valuable resource to nematode research and has the potential to serve as the basis for collaborative improvements in technology for the research community as a whole.

**Fig 1.**
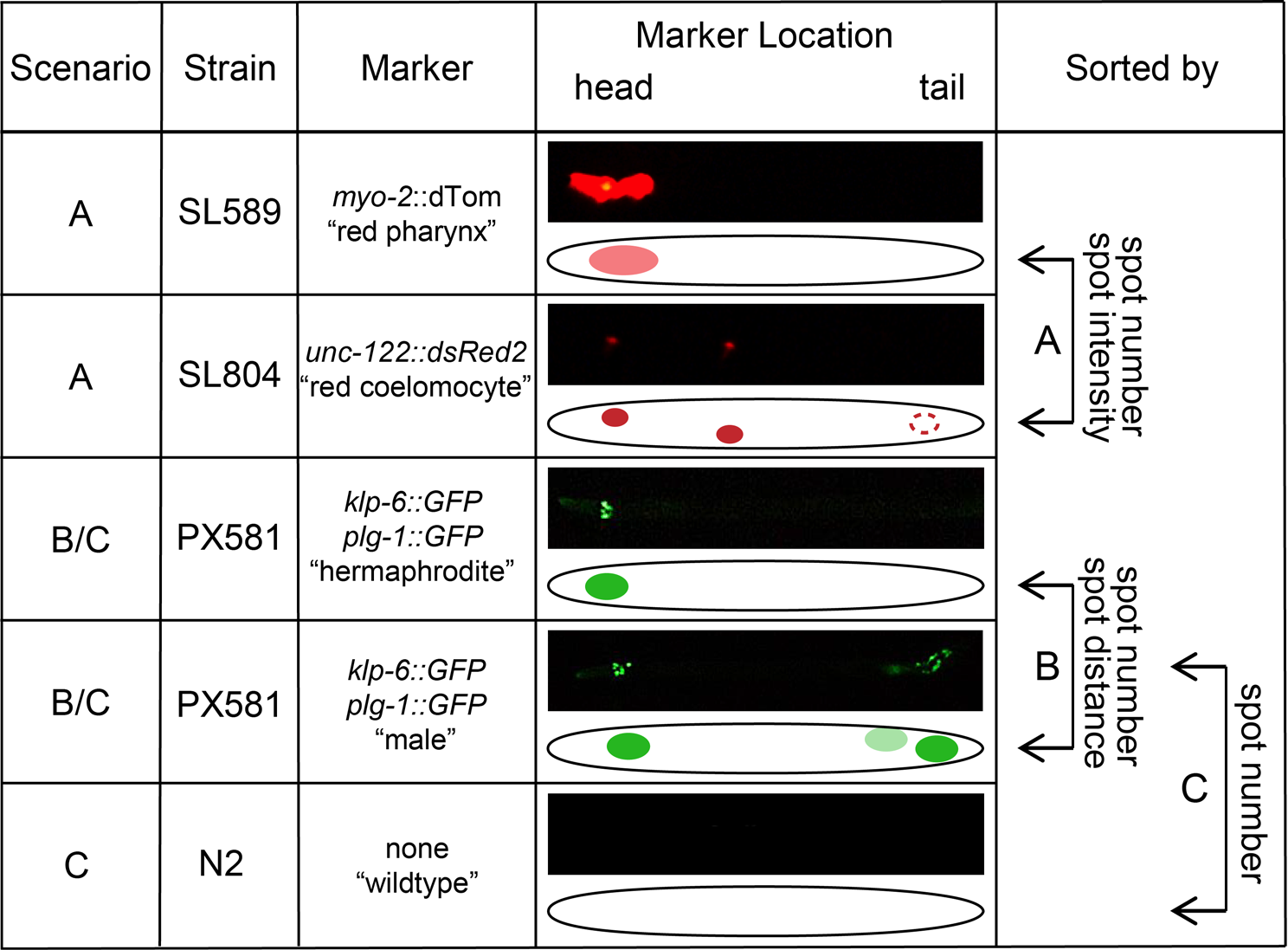
Strain overview. Four different strains were used to test the capabilities of the sorter under three different scenarios (A, B and C). The strains used in scenario A both expressed red fluorescent proteins, one in the pharynx (SL589) and one in the coelomocytes (SL804). For the latter, only the first two pairs of cells are shown, since those were the ones consistently visible during the L4 stage. For scenarios B and C, two other strains are used. Strain PX581 carries two integrated GFP transgenes that make it possible to distinguishing males and females. N2 was used as a strain expressing no markers to be sorted from marked worms. The marker location column shows actual worms in the imaging channel of the sorter. Image contrast was enhanced to show markers more clearly. The sorted by column shows the factors used in automated sorting to determine phenotypic class.

## Methods

### Nematode strains and sorting scenarios

To test the sorter under different realistic Scenarios, we chose three combinations of nematodes (Fig. 1): For Scenario A, we chose two *C. elegans* strains with imaging compatible red fluorescent markers: SL589 (*Pmyo-2∷tdTom*) and SL804 (*Punc-122∷dsRed2*). *myo-2* encodes a pharynx specific myosin, and its well characterized promoter shows strong expression that is restricted to the pharyngeal muscles (strain hereafter referred to as “red pharynx” strain). *unc-122* encodes a transmembrane protein whose endogenous expression is primarily neuronal. Here, we used a well characterized fragment of the *unc-122* promotor [23], whose expression is restricted to the three pairs of coelomocytes, to express dsRed2 (strain hereafter referred to as “red coelomocyte” strain). Although all three pairs of coelomocytes exist in the L4 stage that was sorted, the coelomocyte pairs showed differential expression levels. The observed pattern of expression pattern is most likely a consequence of development as the pairs of coelomocytes located anteriorly and medially originate during embryonic development [24], while the posterior pair arise during larval development [25]. Consistent with the temporal pattern of development, the posterior pair of coelomocytes was reproducibly dimmer than the first two pairs.

For sorting males and hermaphrodites (Scenario B), we generated a strain with sex-specific GFP expression patterns. The strain was generated through standard crosses and carries two independent GFP transgene integrations. The first is a GFP transgene driven by the *plg-1* promoter, a gene that produces a mucin-like protein involved in the formation of the copulatory plug [26]. As expected from the function of *plg-1*, GFP expression is restricted to a spot in the male testis [26]. The second GFP transgene is driven by the *klp-6* promoter. The KLP-6 protein is a kinesin that is expressed in a cluster of neurons in the head of both sexes and in an additional cluster in the tail of males.

For sorting marked and unmarked nematodes (Scenario C), we used the strain from Scenario B (PX581) mixed with age matched wildtype (N2) individuals that do not express any fluorescent markers. For this Scenario, we chose different ratios of marked and unmarked nematodes to determine the relationship between pre-sort ratio and post sort precision. For each sort, we collected two dependent data points: one for the marked and one for the unmarked post-sort precision.

For Scenario C, marked and unmarked nematodes were also sorted without automated strain determination. For these “manual” sorting cases, the experimenter used the live image on the computer screen to make phenotypic determinations for each nematode to be sorted and directed the sorter by keyboard entry of the phenotype. Three populations were sorted using experimenter direction and the speed determined.

### Re-sorting and marker expression optimization

Scenario B was more error prone with respect to detecting the cluster of neurons in the male tail (see Results for details). Misclassification of males results in males being incorrectly sorted into the hermaphrodite exit, therefore reducing hermaphrodite sorting precision. Two approaches were taken to alleviate this problem and maximize final sorting output precision. In the first approach, individuals from the “hermaphrodite” exit tube were sorted a second time. In the second approach the GFP expression in males was optimized for sorting by experimental evolution. This was performed by repeated sorting of the population for more than 30 generations. The male specific posterior *klp-6* spot is typically dim, and only becomes visible late in the L4 state in the original population. By discarding dim males and only keeping the brightest ones to establish the next generation, increased sortability was selected for in this population.

### Fitness consequences of sorting

To determine if sorting has a detrimental effect on individual fitness, a population of nematodes from Scenario B was sorted as described above. Following sorting, 30 single hermaphrodites per treatment were isolated with two males and transferred daily for a week. The total number of off spring produced by each hermaphrodite was taken as a measure of reproductive fitness. As a control, we measured the fitness of 30 unsorted hermaphrodites from the same tube, which had otherwise been treated the same as the sorted individuals.

### Nematode maintenance and preparation

Our sorting chips (Fig. 2 and S1-S3) are sized to sort synchronized L4 nematodes. To generate tightly synchronized populations, axenized embryos were collected by standard hypochlorite treatment and then suspended and rotated in S-basal for 24 hours to generate a synchronized population arrested at the L1 stage [27]. These nematodes were plated at a density of 2000 arrested L1s per 100 mm diameter plate and allowed to develop to the L4 stage. Deviations in population density and developmental timing results in size differences that affect the quality of sorting. The developmental time necessary to reach the optimal L4 stage was established experimentally for every strain used (44, 45, or 46 hours for PX581, N2 or SL589, and SL804 respectively). The arrested L1 nematodes can be plated for four consecutive days, and the L4s can be stored in S-Basal for 3-4 days at 4 °C, without observing body size changes that detract from the quality of sorting. Although extended arrest and exposure to reduced temperature does not alter the ability to sort L4 animals, the altered physiology of such animals should be considered depending on the specific sorter application. For all other nematode handling and processing standard protocols were used [28].

**Fig 2.**
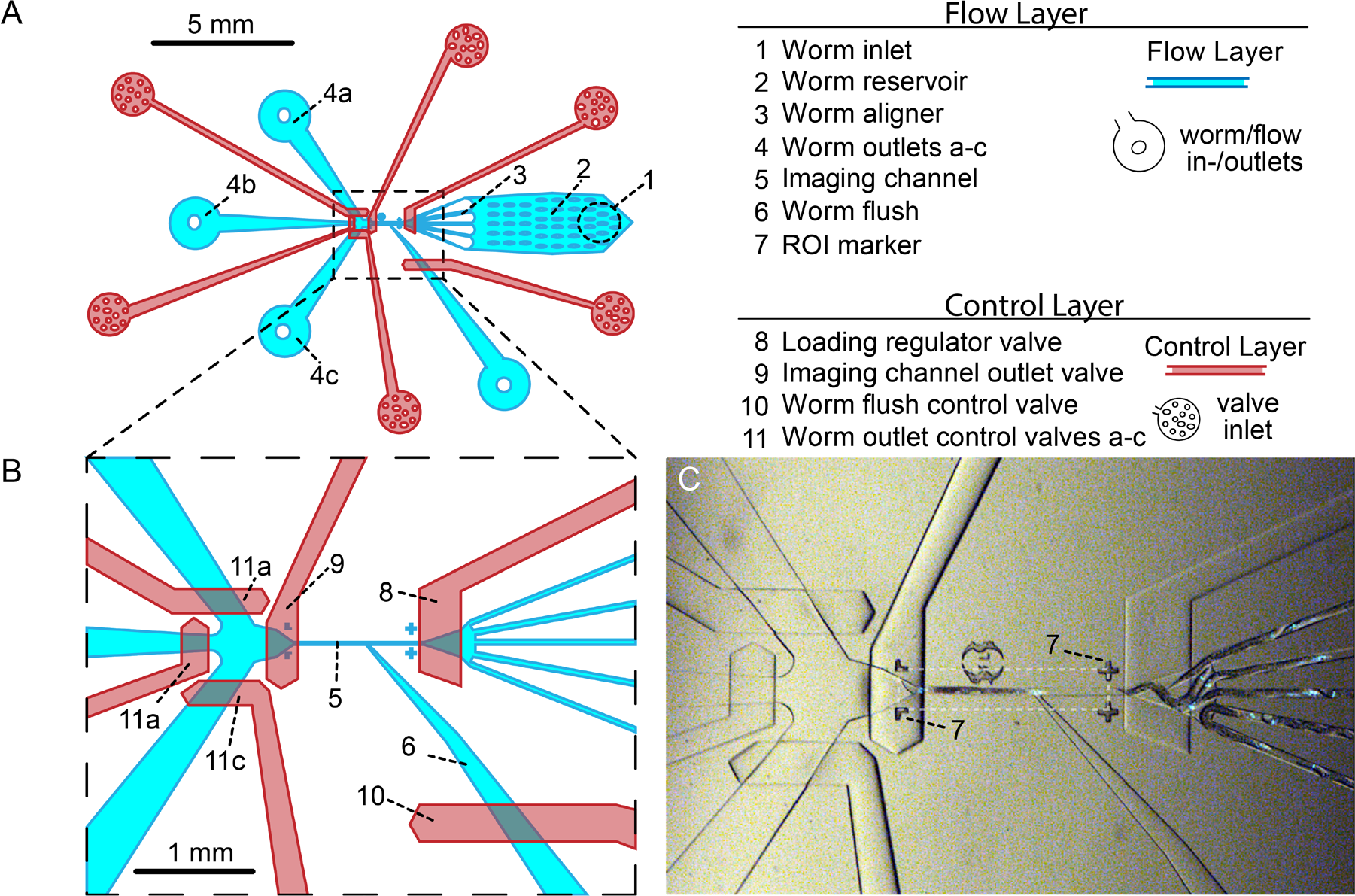
Chip overview. (A) Sorter schematic showing the two-layer design. Worms are directed through the flow layer (blue) by computer-controlled pressurization of the control channels in the control layer (red). In general, worms move from the worm inlet (1), through the worm reservoir (2), and into the worm aligner (3). The worm aligner prepares the worms for entry in the imaging channel (5). (B) When the loading regulator valve (8) opens, a worm can move from the worm aligner (3) into the imaging channel (5). When positioned in the imaging channel an image is captured, processed, and a sorting decision is made. See text for more detailed description. After the decision is made, valve 8 is closed, valves 9 and 10 are opened, and one of the worm outlet valves (11a-c) is opened to allow the worm to be flushed to the appropriate tube. (C) Fluorescent worms in the sorter. The picture shows the same area as the blow-up in B. A PX581 male is positioned in the imaging channel while more worms are prepared for movement into the imaging channel in the worm aligner (feature 3) to the right. The white dashed box shows the typical ROI used for image processing as specified by the ROI markers (7).

For Scenarios A and C, the mixed populations used for sorting were generated by rinsing animals from single genotype populations plates that were grown at 2000 worms per plate. After rinsing the animals in S-Basal, three 10 μl samples were taken from the suspension and worm density was determined by counting L4 animals using a stereomicroscope. The average density of animals in each genotype population was then used to mix the genotypes to the desired ratio for sorting. For Scenario B, the population used was a mixed sex population where reproduction was predominantly via outcrossing. In such a population the sex ratio approaches 50:50.

### Device fabrication

Microfluidic chips were created using standard soft lithography techniques [29,30]. In brief, chip designs were drafted in Vectorworks 2013 Fundamentals (Nemetschek SE, Munich, DE) and photomask transparencies were printed at 20k resolution (CAD/Art Services Inc, Bandon OR, United States). CAD files are available as Supporting Information (S1-S3), and future versions, will be available at https://blogs.uoregon.edu/phillipslabmicrofluidics/. As described above sorters consist of two PDMS layers, one for valve control and one for the nematodes to flow though. Following the procedure for standard photolithography, the design for the two layers were printed on separate photomask transparencies, which were then used to pattern two SU-8 photoresist masters (see S4 for more details).

These masters served as molds. To make the thinner control layer, a thin layer (~5 g) of PDMS (Sylgard, PDMS:developer ratio 1:20) is applied onto the flow layer mold by spin coating and partially cured by incubating for 28 min at 60 °C. Spin coating results in a thin layer of PDMS covering the channels, which ultimately becomes the septum between the flow and control channels in the finished device. The partially cured layer is slightly tacky and will stick to the naked finger for 1-2 seconds when at the right degree of polymerization. To make the thicker control layer, approximately 20 g of PDMS (PDMS:developer ratio 1:5) is poured onto the control layer master and incubated for 30 min at 60 °C. After cooling to room temperature, the control layer is cut out and aligned with the flow layer, which is still attached to the mold. After alignment, both layers are incubated together for 1.5 h at 60 °C to fully cure. After cooling to room temperature, the individual sorters are cut out and holes for connecting tubing to the control valve (Fig. 2, valve inlets) and flow (Fig. 2, worm/flow in-/outlets) channels are punched. While all fluid connections are made with 1.5 mm stainless steel tubes, only the flow in/-outlets are punched at 1.5 mm. To maintain the tubing-PDMS connection during the periodic pressurization to 30 psi, the valve inlets were punched with a 1 mm disposable biopsy punch to generate undersized holes that more tightly fit the 1.5 mm OD stainless steel tubing. When the valves are pressurized, small tears or micro-fissures in the PDMS at the inlet-stainless steel tubing interface can cause connection failure. Because dull biopsy punches increased the frequency of micro-fissures, and therefore increased the rate of connection failure, the usage of the biopsy punches was tracked, and punches were discarded following a maximum of 150 uses. Following these manipulations, the devices are plasma bonded to a glass slide, which serves as the floor of the flow layer. After a final bake at 60 °C for 1.5 hours, the device is ready to use.

### Control system

In addition to the PDMS microfluidic chip, a functional sorter setup requires a control system that is linked to a computer running the necessary software (see *Image acquisition, processing and sorter control*). The sorter control system consists of a set of regulators, valve-controllers, and a compressed air source (Fig. 3, a list of all parts is available as S5). The input air is set to 40 psi by the first regulator and then split up into two tubes. Using additional regulators, one stream of air is set at ~10 psi (can be between 5 and 15 psi depending on worm size and desired speed) and divided using a T-junction to feed into two 15 ml conical tubes: one for the flush and one for the buffer containing the nematodes. The tubing enters the 15 ml conical tube and terminates directly beneath the lid such that the incoming air pressurizes the air above the liquid, ultimately forcing the fluid out of the 15 ml conical tube via a second piece of tubing whose end sits just above the bottom of the tube (see Fig. 3).

**Fig 3.**
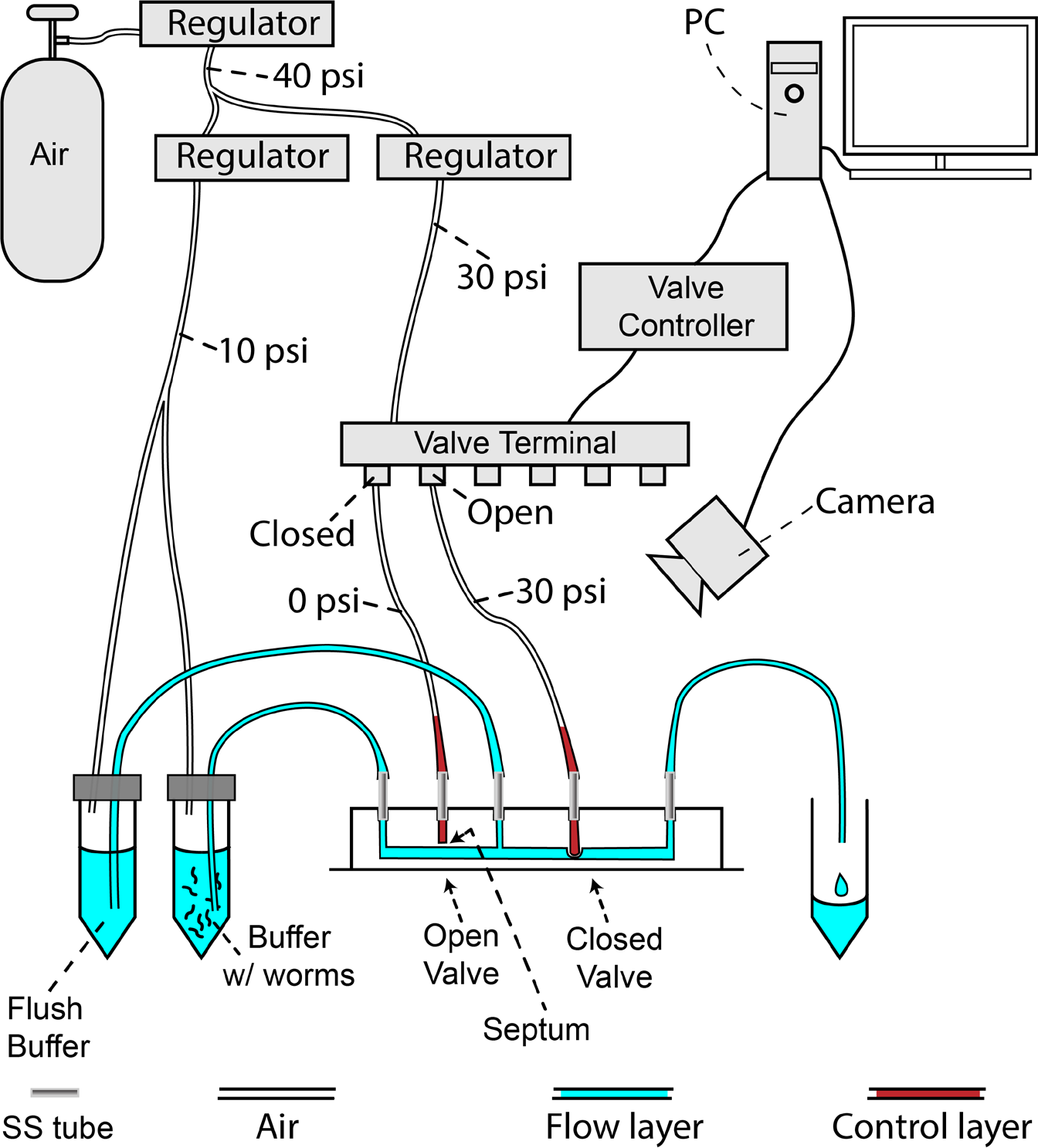
Schematic sorter overview. The fluid in both layers of the sorter are pressure controlled. The flow layer is pressurized constantly during a sorting run (~10 psi), while the control layer can alternate between 0 and ~30 psi. The pressure state of the control layer is determined by the MATLAB code, which controls the open/closed state of the valves in the valve terminal (Festo, S5) via the valve controller (WAGO, S5). When a valve in the valve terminal opens it pressurizes a control layer channel, causing the septum dividing the control layer channel from its associated flow layer channel to distend into the flow layer. This distension acts as a microfluidic valve, blocking flow at that point in the flow layer.

The second 40 psi air stream is down regulated to 30 psi using a third regulator and connected to the air-valve array (FESTO, S5) that opens and closes individual mechanical valves, ultimately controlling the control valves on the chip. This array is controlled by the valve controller (WAGO, S5), whose actions are in turn determined by the software running on the computer. In addition, the computer processes images obtained via a microscope camera and determines which channel to sort the individual nematode into. In the example in Fig. 3, only two control channels are connected to the air valve controller, with the left one closed and the right one open. On the chip (shown as a lateral cross section), this leads to the control channel being pressurized and the septum expanding into the flow channel underneath. This expansion is possible because of the elasticity of the very thin PDMS septum that separates the control channels from the flow channels. Once the valve is activated, the flow in the flow layer is stopped, and nematodes are not able to pass though the chip at that location. When the pressure is released the PDMS septum rebounds, and the nematodes can be flushed though the channel into the outlet.

### Device setup

The device is placed under a stereo microscope (Leica M205 FA) equipped with a computer-controlled camera (Leica DFC 310 FX). The initial step in setting up the sorter is to connect the valving tubes to the valving channels on the chip in a predefined order (see S4). One end of the tubing is equipped with a 1.5 mm OD stainless steel tube that serves as a connector. This end is filled with 2 cm of water and connected to the holes punched into the valving channels. Inserting the metal connector requires some practice to avoid ripping the PDMS. When the valve tubing is successfully inserted the other end of each tube is connected to the valving controller via Leuer connections. After all tubes have been inserted, the valves are pressured to 20 psi using compressed air. The water will slowly displace the air in the valving channels because PDMS is permeable to air but not water. Filling the valves with water has the advantage that no air is driven into the flow layer when the chip is operational. The filling process is then observed through the microscope until all of the valves are filled (~10-20 min), at which point the pressurized air is turned off. While filling the valves (or thereafter) the remaining inlet, outlet, and flush tubes are connected, and the nematodes can be flushed in. The detailed protocol for setting up a sorter is accessible as Supporting Information (S4).

### Image acquisition, processing and sorter control

The sorter is controlled by a MATLAB program (code for the three scenarios: S6-S8) that directs the pressurization of the on-chip valves, as well as coordinating with an external image-acquisition program (μManager [31,32] in our case) to acquire and process the nematode images. During device start-up, lighting, focus and magnification (90x) need to be optimized and valve filling and control initiated. At this stage, the device is ready for user directed sorting, or automated sorting can be initiated. If automated sorting is initiated, the experimenter will be presented with a sample screen image and instructed to specify the upper left and lower right of the target imaging zone (Fig. 2, feature 7). The software will then prompt the experimenter for the number of nematodes to be sorted, after which sorting begins automatically.

For automated sorting, any camera software capable of displaying a live view can be used. The user will specify a region of interest using the supplied MATLAB code, and an integrated JavaScript screen capture script is used to capture an image of each individual passing through the imaging channel (see ROI Fig. 2C) which is then processed as illustrated in Fig. 4. Fig. 4A shows the nematode in the channel with bright field illumination. When automatically sorting fluorescently marked individuals, no white light is used. The software determines each worm’s phenotypic class by analyzing the distribution of fluorescence within the image by: (1) converting the image from grayscale to a black and white using a preset threshold (Fig. 4B), (2) expanding the edges of GFP positive puncta using a MATLAB blurring function (without tapering) to connect closely located GFP positive cells (Fig. 4C), and (3) using the criteria specific to a given sorting Scenario to categorize the phenotype of each individual based on the number and distribution of fluorescent spots (Fig. 1 and Fig. 4). Any divergence from defined parameters for the sorted Scenarios leads to flushing into the waste outlet, since it may indicate the presence of two or more individuals in the imaging channel, which would compromise the precision of the sorting.

**Fig 4.**
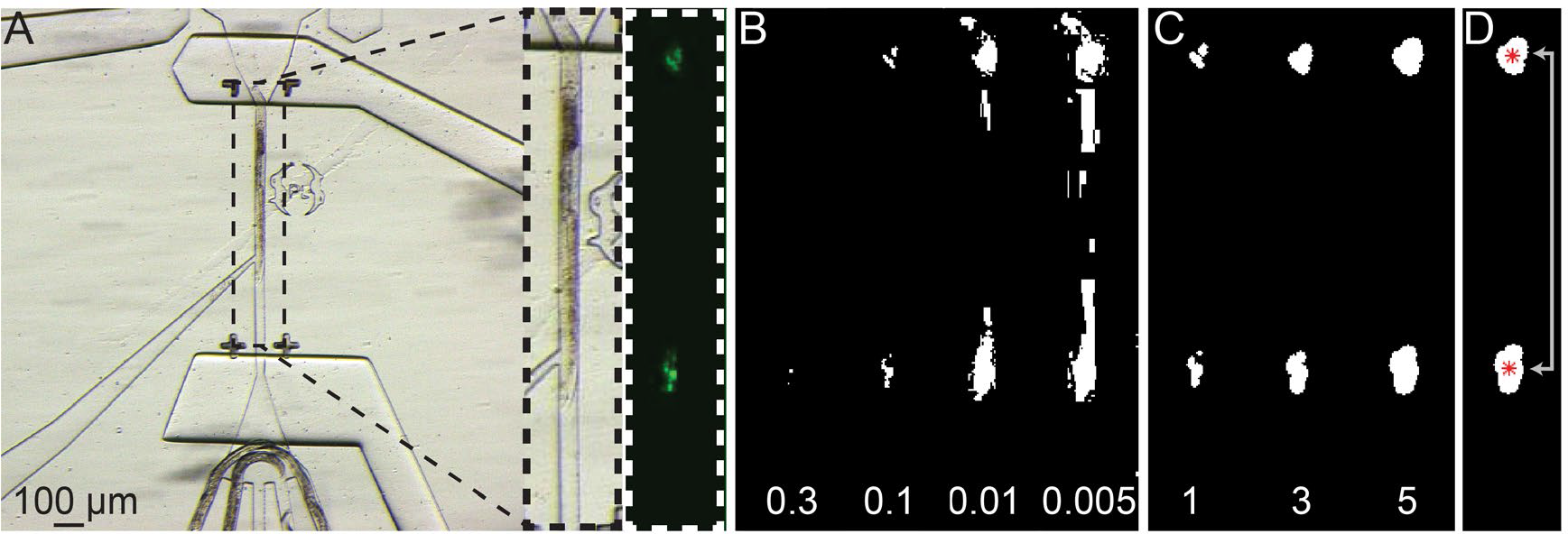
Image processing workflow. (A) Brightfield image of a worm in the imaging channel, the selected region of interest, and the GFP image to be processed by the custom MATLAB code (S6-8). (B) The image is first converted to a black and white image. Shown are four converted images using a range of thresholds from 0.3-0.005. (C) Images are then processed to connect nearby white pixels. Shown is the image processed with a black/white threshold of 0.1, with connection parameters of 1, 3, and 5. (D) Centroids for each contiguous patch of white pixels is determined, centroid number is counted, and distance between the two furthest centroids is measured.

After the phenotype has been determined, the nematode is flushed into the appropriate outlet by opening the imaging channel outlet valve, flush valve and the relevant control valve (Fig. 2b, features 9, 10, and 11a-c). A video of nematodes from Scenario B can be found as Supporting Information (S9). Upon completion of a sorting experiment, nematodes still remaining in the device are flushed into the appropriate exit tubes by opening all valves (Fig. 2b, features 8-11), disconnecting the inlet port (Fig. 2a, feature 1) from the conical with buffer and worms (Fig. 3), and allowing a continuous flow of buffer to come from the flush reservoir. After flushing the worms from the channels and the nematode reservoir, the chip can be reloaded for additional sorting runs.

### Data acquisition

Sorting precision under the different Scenarios was determined by sorting multiple replicates of independent test populations, with each replicate being tested on a different day. For every sorting replicate, the precision, *Q*, was calculated for both sorted subpopulations after sorting to their respective outlets. The precision for each outlet was calculated as

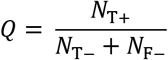

where *N*_*T*+_ is the number of true positives, or correctly sorted individuals, and *N*_*F*−_ is the number of false positive, or incorrectly sorted individuals, for that outlet. Five replicates for Scenario A, six for B and eleven for C were processed. These replicates are presented in chronological order and no sorting replicate was excluded from graphical representation or statistical analysis.

The nematodes where prepared as described above and each sorter run was conducted for two to three hours. After the nematodes were collected from the outlet, 0.5–1 ml of S-basal supplemented with 0.01% Triton X-100 was added to keep nematodes from sticking to tube and pipette tip walls. Nematodes were allowed to settle in a tube to reduce the volume and then were plated onto seeded NGM. The nematodes were allowed to recover overnight and were phenotyped and counted the next morning.

### Data Fitting

To characterize the relationship between the frequency of a phenotypic class in the initial population and the resultant sorting precision we derived equations relating the observed precision to the properties of the sorting process (see Scenario C: Marked vs. Unmarked below). Using the derived equations and the observed precision/phenotypic frequency data we fit the data to solve for the underlying sorting accuracies. Data-fitting was performed using a non-linear least squares method with bisquare weighting and the default trust-region-reflective algorithm in the MATLAB curve fitting app. The starting point for the two accuracies was set equal to random sorting (0.5), and the bounds were set to zero and one.

### Material costs

The costs for a sorter can be split into one-time hardware costs and costs for the actual chip fabrication. The control hardware without the microscope and the computer running the software generates costs of about $2000 US. Making a set of two reusable masters costs approximately $180 US if a clean lab with photolithography, equipment spin coater, programmable hotplate and plasma cleaner are available. The marginal cost for production of one chip is approximately $2.50 US. Additional potential startup costs would include a spin coater, plasma cleaner, and low-temp oven, if a facility for microfluidic fabrication is not already available. A full list of materials used is accessible as Supporting Information (S5).

## Results

### Microfluidic design

To direct nematode movement through the flow layer of the device, sorting chips have a control layer bonded on top of a flow (sample) layer [33]. In our design, there are six independent control channels in the control layer for which pressurization causes the septum separating the control channel from its specific flow channel to distend into and block the flow layer wherever the two layers’ channels cross (see device fabrication for more details).

The chip design presented here (Fig. 2) is derived from the previously published design by Crane et al. [5]. In addition to numerous minor modifications, three additional features were added. First, one more exit (Fig. 2a feature 4b) and associated control valve (Fig. 2b, feature 11b) were added. Second, while prototyping sorter designs, it proved necessary to add another valve (Fig. 2b, feature 8) to the entrance of the imaging channel as a loading regulator [21] to prevent multiple individuals from entering the imaging channel (Fig. 2b, feature 5). Third, a flush channel (Fig. 2b, feature 6) was added that conveys the individual from the imaging channel to the exits while minimizing the potential for simultaneously dragging along additional individuals from the worm aligner (Fig. 2a, feature 3).

The function of each of these features becomes clear when following an individual though the chip (Fig. 2): when an individual enters into the device, it enters through a 1.5 mm worm inlet (feature 1) that is punched in the arena with pillars (feature 2). The arena with pillars acts as a small nematode reservoir to ensure consistent animal loading into the imaging channel. To facilitate quick, single entry into the narrow imaging channel, the nematodes flow through one of the five channels connecting the reservoir to the imaging channel. These channels align them (feature 3) in preparation for movement into the imaging channel (feature 5). Movement from the worm aligner into the imaging channel is controlled by the loading regulator valve (feature 8). If the valve is open, an individual can enter the imaging channel where their progress is blocked by the next valve (feature 9) that controls exit from the imaging channel. While in the imaging channel each individual is phenotyped either by the experimenter or by automated image analysis. After a sorting decision is made, the loading regulator valve behind the individual is closed. Three valves then open to flush the individual into the correct outlet: the valve that previously blocked exit from the imaging channel (feature 9), the valve that controls the flush channel behind the individual (feature 10), and the valve controlling the selected outlet (feature 11). The buffer coming from the flush is responsible for carrying the nematode into the outlet.

If the individual is not successfully flushed at this point, then there is the possibility to re-flush via repeating same procedure. After this cycle, the valves switch back to the original configuration, thereby allowing the next individual to enter the imaging channel (3D animation of sorter functionality can be found as Supporting Information (S10)). In principle, any number of outlet channels could be added to the system for more complex sorting arrangements, although there will be space limitations for the valves on the control layer.

### Scenario A: Marked vs. Marked

After sorting a mixed population of red pharynx and red coelomocyte nematodes (50:50), the red pharynx collection tube contained 97 ± 2% pharyngeally marked individuals and the red coelomocyte collection tube contained 95 ± 2% coelomocyte marked individuals (Fig. 5). The overall yield, meaning how many individuals were sorted into the exits relative to the waste exit, was about 63 ± 6% for this scenario, indicating that a substantial portion of individuals could not be categorized by the software. Sorting speed (measured over three independent replicates) was relatively high with an average of 1426 ± 230 individuals/hour. The observed yield ranged between 625 ± 35 animals/hour (red pharynx) and 335 ± 109 animals/hour (red coelomocyte).

**Fig 5.**
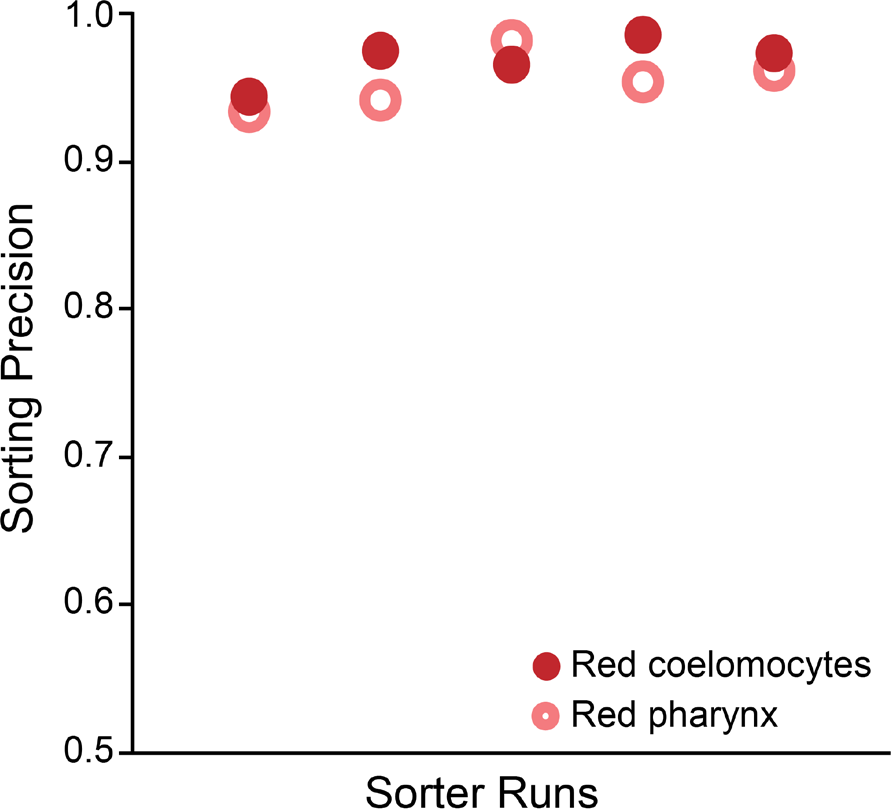
Sorter precision for differentially marked animals (Scenario A). The five replicates for Scenario A exhibit qualitatively indistinguishable after sort precision (range 0.93-0.98, Wilcoxon *p*=0.21). Also, the sorting precision of the two strains does not differ within the runs (with a maximum inter-strain precision difference of 0.03).

### Scenario B: Males vs. Hermaphrodites

Both males and hermaphrodite were enriched from an approximate 50:50 sex ratio to 81 ± 6% hermaphrodites in the hermaphrodite outlet and 92 ± 2% males in the male outlet. The male precision was usually substantially higher than the hermaphrodite precision (Fig. 6). The overall yield (number of individuals sorted into the nematode exits versus the waste) was about 75 ± 15 % for this scenario, and speed was 774 ± 202 nematodes/hour.

**Fig 6.**
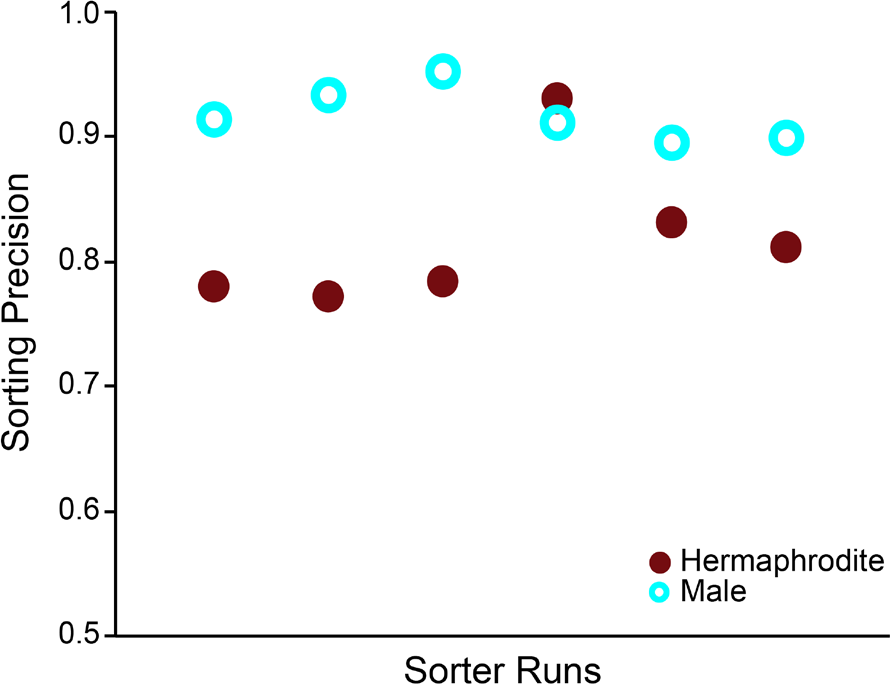
Sorter precision for hermaphrodites vs. males (Scenario B). Five of the six replicates show the same pattern in the sorting precision, with the precision of male sorting being higher than the that for hermaphrodites (Wilcoxon *p*=0.03), resulting from a better detection of females than males by the software, meaning that males were more likely to be missorted. The fourth sort was done with slightly older worms that had to be sorted slower due to their larger size. The improved precision for that replicate could be the result of stronger marker expression due to the older age, the decreased speed of sorting due to increased size, or both.

### Re-sorting and marker expression optimization

A single additional round of re-sorting the hermaphrodite population helped to increase hermaphrodite precision to the level of the males (93%). Additionally, two replicates of the sorting-adapted population were sorted. On average, both males and hermaphrodites from the evolved line had an after-sorting precision of 92 ± 2%. Thus, precision in sorting of hermaphrodites was greatly increased after selecting for stronger expression of the male-specific markers.

### Fitness consequences of sorting

To determine if sorting had a detrimental effect on individual fitness, the brood size for sorted and non-sorted individuals was measured (see materials and methods). The fitness of sorted and non-sorted individuals did not significantly differ from one another (Fig. 7; *Wilcoxon ranked sums* test, *Z* = 0.46, *df* = 1, *p* = 0.649).

**Fig 7.**
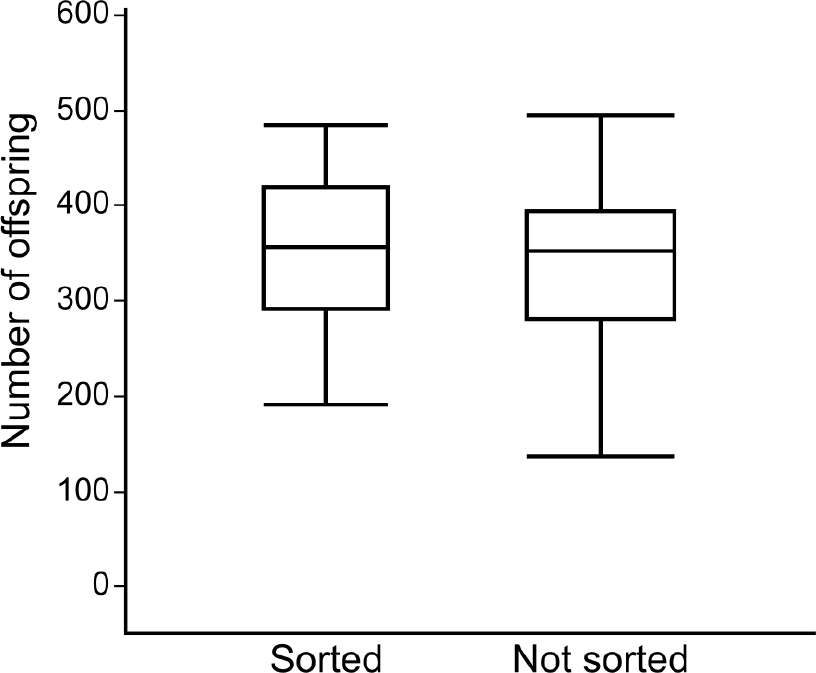
Potential consequences of sorting on offspring production. There is no significant difference in fitness, measured as the total number of offspring, between sorted and unsorted worms (Wilcoxon *p*=0.649).

### Scenario C: Marked vs. Unmarked

To analyze the relationship between the frequency of a phenotypic class in the starting population and sorting precision, a series of experiments with different ratios of marked and unmarked individuals in the starting populations were conducted. For these experiments the overall yield was 100% since our sorter cannot detect unmarked individuals in this configuration and will not send any nematodes to the waste outlet. Any error in sorting will be directly reflected in the precision of the resulting populations. Marked and unmarked individuals were separated at a rate of 1000 ± 289 nematodes/hour. The observed precision of the sorted subpopulations appeared dependent on the frequency of the phenotype in the starting population (Fig. 8). To explore this dependency, we considered the relationship between the accuracy of sorting individual animals and the ultimate precision of the sorted subpopulation. To do so we first defined the relevant properties of the starting population that is to be sorted. We define *f* as the frequency of individuals (worms) in the starting population that are true positives. It is also the probability that an individual presented to the sorting system is a true positive. Conversely, the probability that an individual presented to the sorting system is non-positive is (1 − *f*).

**Fig 8.**
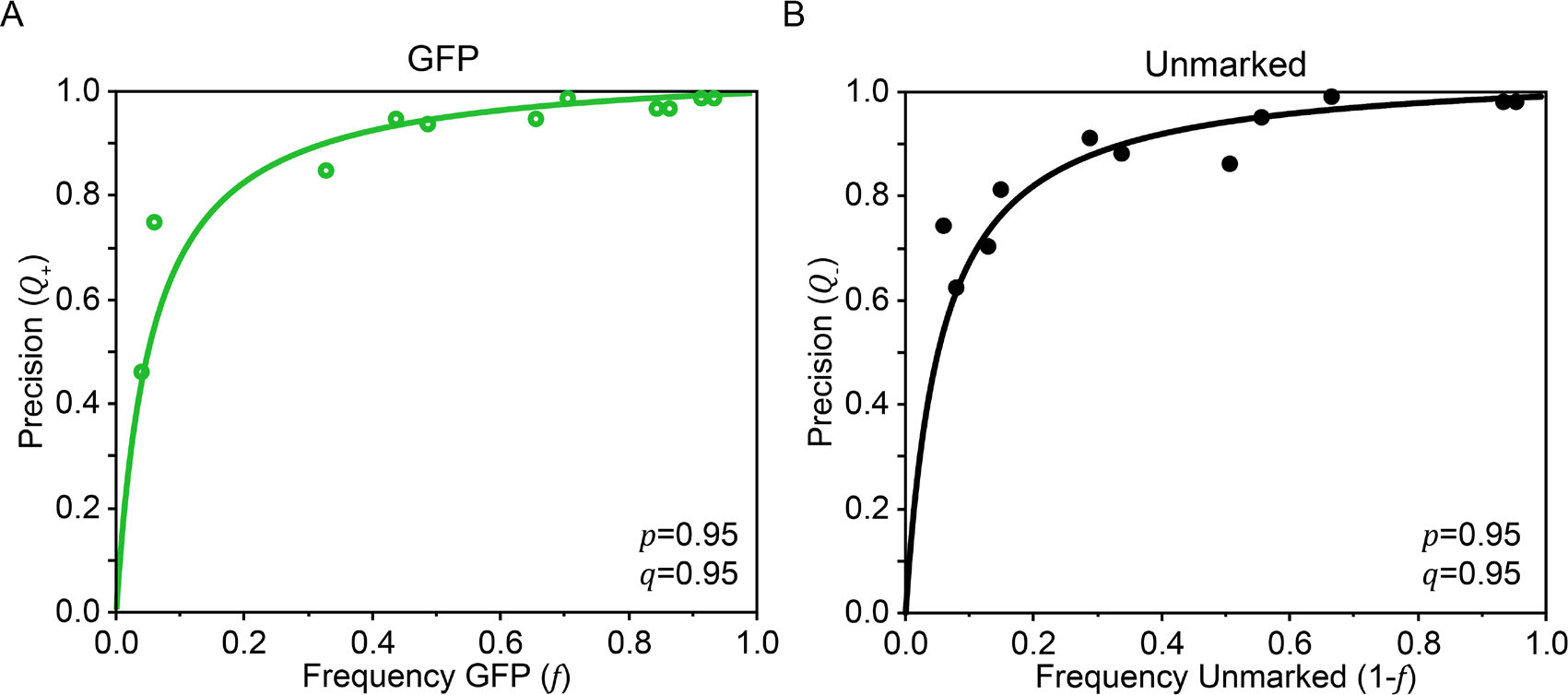
Sorter precision for marked vs. unmarked worms (Scenario C). The precision, Q, for each sorted GFP marked (A) and unmarked (B) subpopulation is shown as a function of that subpopulations frequency in the starting population (dots). The solid lines show the calculated precision using equations [1] and [3], and the values for *p* and *q* obtained by fitting the observed data. For the GFP marked data (A) the goodness of fit statistics are *SSE*= 0.0121, adjusted *R-square*=0.948, and *RMSE*=0.03667. For the unmarked data (B) the goodness of fit statistics are *SSE*= 0.002196, adjusted R-square=0.8442, and *RMSE*=0.0494.

We then defined the relevant sorting system properties. We define *p* as the probability that the sorting system will call a truly positive individual positive, and *q* as the probability of the sorting system calling a truly non-positive individual non-positive. Our performance assessment of the sorting system, “precision”, can be understood in the context of the properties of the population and the sorter system. Precision, *Q*, measures the ability of the sorting system to correctly classify true positives (T+) as positives or true negatives (N–) as negatives. For example, the precision of sorting true positives, *Q*_+_, is defined as the fraction of true positives in the subpopulation sorted as positive by the sorting system. In terms of population properties and sorting system properties, this is

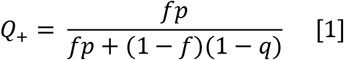

Each term in this expression is the joint probability of two events. The first term, *fp*, it is the probability of picking a true positive to be sorted (*f*), *and* sorting it correctly (*p*). In other words, this represents the probability of an individual in the starting population completing being sorted as a true positive. The second term, (1 − *f*)(1 − *q*), is the probability of picking a true negative to be sorted (1 − *f*), *and* sorting it incorrectly (1 − *q*). This represents the probability of an individual in the starting population being sorted as a false positive. Multiplying the top and bottom of the right hand side of this equation by *N*, the total number of individuals sorted, provides a computational formula that can be applied to experimental data,

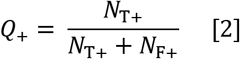

where *N*_*T*+_ and *N*_*F*+_ are the number of true positives and false positives, respectively. There are analogous expressions for measuring the ability of the sorting system to correctly classify true negatives. They are,

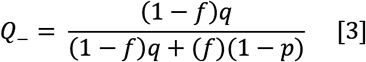

and

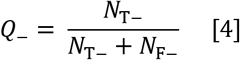

The sorting system properties *p* and *q* can be estimated by plotting experimental values of *Q*_+_ or *Q*_−_ equations [2] and [4], against *f* and fitting the data with equations [1] and [3], respectively. Doing so, we obtained the values of *p* and *q* used to generate the fitted curves in Fig. 8. These values can be interpreted as the accuracies of the sorting system for true positives (*p*) and true negatives (*q*). The curves generated using *p* and *q* in equations [1] and [3] illustrate the variation in sorting precision that can be generated regardless of a fixed sorting accuracy. For example, despite a high sorting accuracy for GFP positive individuals (*p* = 0.95), the precision drops precipitously when the frequency of the GFP positive individuals drops below 0.2 (Fig. 8a). Multiple sorting runs will probably be necessary for low-frequency markers and/or screens for rare individuals.

To compare automated image processing with user determination of phenotype, we sorted nematodes in the microfluidic device using the optional keyboard input. We performed three sorter runs with user input, each for sorting durations longer than 60 min. Marked and unmarked individuals were sorted at an average speed of 1260 nematodes/hour. False sorting was very rare and could be directly seen while sorting. Nevertheless, a subset of one of the runs was scored afterwards with only one unmarked worm out of 300 worms (precision = 99.7%).

## Discussion

Using our sorter platform, we were able to perform automated sorting of nematode populations with repeatable results. Our sorter platform is open source, relatively inexpensive, and independent of imaging platform used. We find that sorting precision is dependent upon the scenario and the starting ratio of marker types. Depending on the sorting scenario, the rate of nematode sorting ranged from 774 to 1426 nematodes per hour, a rate that falls in the middle of previously published microfluidic sorters. For example, Crane et al. [5] were able to sort worms at a higher speed of 1500 to 2500 worms/hour using a design that is simpler than ours, but did not report the sorting precision of their devices. On the other end of the spectrum, Chung et al. [21] tested two sorting scenarios with a marker ratio similar to our Scenario A (~30:70 vs 50:50) in their device. Their sorting speed was 150 to 400 worms/hour depending on their scenario; about 1/8 to 1/3 the speed that we achieve here. They report a precision that is comparable to our precision from Scenario A (0.91 to 0.98 compared to 0.95 to 0.97) (Fig. 5). Even so, microfluidic chips cannot compete speed-wise with commercially available flow-cytometry based sorters. For example, Union Biometrica reports the maximum speed for their COPAS as 25 events/s, or 90,000 events/hour [34].

Increases to speed and precision should be possible via further optimization of the automated sorting algorithms and chip designs. Additionally, the increased sorting precision after experimental evolution of brighter *Pklp-6∷GFP* expression suggests that judicious choice of markers should greatly increase sorting precision, as well as speed.

## Conclusion

Using a nematode sorter can make many experiments easier to perform at a scale that is more likely to provide meaningful results. We were able to differentially sort individuals with high precision under three different scenarios, covering a range of potential applications. Nevertheless, further developments in this approach will be necessary to match existing (very expensive) commercial solutions. Together with other nematode applications for microfluidic chips, this technology represents a reasonable, powerful and very flexible tool for nematode research.

## Supporting information

S1 Sorter File_VWX

S2 Sorter File_DWG

S3 Sorter File_EPS

S4 Sorter Protocol

S5 Parts List

S6 Code Scenario A

S7 Code Scenario B

S8 Code Scenario C

S9 Sorting Video Scenario B

S10 Sorter Animation

## Acknowledgements

The authors would like to thank B. O’Hagan, M. Rockman and the *Caenorhabditis* Genetics Center for providing nematode strains. We also thank Christine O’Connor for statistical assistance, Benjamin W. Blue for providing the 3D animation, Mario Ranallo for technical assistance in the manufacture of the chips, and Kurt Langworthy and the Lokey Labs nanofabrication facility for access to fabrication equipment. Conversations and schematics from Hang Lu and her lab were critical for helping us get started with our initial designs.

## Funding

This work was supported by NIA grants R01 AG049396 and R21 AG043988 to PCP.

## Accessibility

All protocols (S4), MATLAB code (S6-8), data (S11), and design files (S1-3) are available in the Supporting Information. Current and future sorter designs will also be made available on our microfluidics webpage (http://blogs.uoregon.edu/phillipslabmicrofluidics).

## Supporting Information

**S1: Chip design.vwx** Sorter design as Vectorworks file.

**S2: Chip design.dwg C+F** Sorter design as dwg. One for the control and one for the flow layer.

**S3: Chip design.eps C+F.** Sorter design as Encapsulated Postscript file. One for the control and one for the flow layer.

**S4: Protocol device Fabrication and Sorter Setup.** Detailed protocol on how to manufacture masters and chips and how to set up a device for sorting.

**S5: List of Parts Used.** Parts needed for making a master, a set of sorter chips and the actual sorter are listed together with costs and supplier information for the USA.

**S6: Code Scenario A.** MATLAB sorting code for separating labeled pharynx and coelomocytes nematodes from each other.

**S7: Code Scenario B.** MATLAB sorting code for separating labeled hermaphrodite and male nematodes from each other.

**S8: Code Scenario C.** MATLAB sorting code for separating labeled and unlabeled nematodes from each other.

**S9: Sorting Video Scenario B.** Video showing sorting hermaphrodites from males. This video shows a sequence sorted by hand under bright light plus GFP settings. Automatic sorting requires only GFP setting without white light.

**S10: Sorter animation.** 3D animation of the sorter design and functionality.

**S11: Data.**

