## Supplementary material for "An open source microfluidic sorter for *Caenorhabditis* nematodes": S4 Sorter Protocol

**Phillips Lab**

**Microfluidics Protocol: Sorter**

**Master fabrication**

A) Wafer Cleaning

*Wafers come in different qualities and levels of cleanliness. The following method is for wafers with stains and particles attached to the surface. Rinse cleaning can be reduced or skipped if wafers come clean out of the box. Plasma cleaning should always be used.*

**Rinse Cleaning**

1. Rinse ethanol over the wafer
2. With gloved finger rub the surface of the wafer
3. Rinse the wafer with isopropyl alcohol (IPA)
4. With a different gloved finger, rub the surface of the wafer
5. Rinse the wafer with IPA
6. Dry wafer using house air and then place wafer in petri dish

**Plasma Cleaning**

1. Place wafer in plasma cleaner (Harrick Plasma) and vacuum the chamber
   - After vacuum the chamber for ~2min open the chamber to the needle valve
   - Turn on the plasma for 30 seconds
   - Turn off plasma and open the chamber to ambient air
2. Place wafer back into a petri dish

B) Applying Photoresist

*Sorters need two masters, one for the flow and one for the control layer. The manufacturing for these differs only in the spinning speed needed to create wafers of different heights. The spin coater holds the wafer in place by vacuum. The photoresist is very viscous and using a black 50 ml conical tube is advisable for easier portioning. Fill the tube ahead of time to let air bubbles have time to disappear.*

**Prepare Spin Coater** (Headway Research, Inc.)

1. Set 2 step program
   - Step 1 (distribution of photoresist): 10 sec, 500 RPM, 100RMP/sec
   - Step 2 (spinning to height): 30 sec, 300RMP/sec, RPM according to table

| **Layer** | **Feature height** | **Photoresist** | **RMP Step 2** |
| --- | --- | --- | --- |
| flow | 35 um | SU8 2025 | 2250 |
| control | 15 um | SU8 2025 | 4500 |

2. Place paper/foil around the spinner for ease of cleaning
3. Place proper chuck into the device
4. Open vacuum valve
5. Place a clean wafer onto the chuck, turn on the vacuum and test run the procedure

**Spinning on SU-8**

1. Place wafer onto the center of the spin coater
2. Turn on the vacuum
3. Pour Hershey kiss amount (2.5 ml) of the proper SU-8 onto the center of the wafer and cover the spin coater with lid
4. Start Program
5. Once the program has finished running turn off the vacuum
6. Place wafer back into the plastic petri carrying dish
7. Repeat steps 3-6 for each wafer
8. Clean the spin coater
   - Remove the foil/paper and throw away
   - Use acetone to wash down the inside of the spin coater
   - Use acetone to wash the chuck
9. Close the vacuum pipe
   - Turn off the spin coater
10. Refill the SU-8 tube that was used

**Soft Bake (Torrey Pines Scientific)**

1. Program (temp, time, ramp):
   - 65 C, 4 min, max
   - 96 C, 8 min, max
   - 45 C, 1 sec, max
2. Starting the program
   - Place wafers onto the hotplate
   - Cover each wafer with a glass petri dish
   - Start the program
   - Cover all of the glass petri glasses with aluminum foil
3. When the hot plate has cooled bellow 40^o^C it can be removed from the hot plate and placed into a plastic petri dish

C) Exposure

*Wear appropriate eye protection when exposing as UV light can damage eyes*

1. Turn on light source (UV lamp), pre-heating might be necessary
2. Place metal mask holder onto the mask aligner and attach the hose to house vacuum (the mask aligner is too small in our facility)
3. Turn on timer and then set timer to 4 seconds
4. Check the power on the UV illuminator by placing the light probe onto the center of the mask holder and exposing (Moving the stage under the light source)
5. Use the light probe reading to determine the appropriate exposure time (Table in the appendix with reference exposure times)
6. Set the exposure time on the mask aligner to the previously determined time
7. Place wafer onto the circle in the center of the metal mask aligner stand
   - Point the flat edge of the wafer towards the top of the mask holder stand
8. Place emulsion mask onto the wafer with the emulsion side faced down, align the mask so it covers the wafer with the mask description on the flat side (generally place the mask with the mask name readable)
9. Turn on the vacuum (securing the mask onto the wafer)
10. Move stand under the UV light (**look away even when wearing glasses**)
    - The UV light will automatically turn on and expose for the proper time and then turn off
11. After exposure move the stand out from the light
    - Turn off the vacuum
    - Remove the mask from the wafer (wafer may slightly stick so remove slowly and gently)
12. Move the wafer back to the hotplate
    - Cover with glass petri dish
    - Cover with Aluminum Foil
13. Repeat steps 7-12 for each wafer
14. Turn off the Mask Aligner
    - Turn off the UV light source
    - Turn off the Mask Aligner timer
    - Remove the metal mask holder and reconnect the mask aligner vacuum house (specific to our type of mask aligner)

**Post Exposure Bake (hot plate)**

1. Program (temp, time, ramp):
   - 65 C, 1 min, max
   - 96 C, 8 min, max
   - 30 C, 1 sec, 120 C/h
2. Set and start hotplate program
3. When the hot plate has cooled bellow 40^o^C, the wafer can be removed from the hot plate and placed into a plastic petri dish

D) Developing

1. Submersion Development for 10 minutes
   - Place wafers into the PMEA container rack
   - Close the container
   - Place the container under hood
2. Remove wafer from PMEA
3. Rinse both sides of the wafer with PMEA into chemical waste
4. Rinse both sides of the wafer with Isopropanol into chemical waste
5. Dry wafer using N2 or house gas
6. If there are streaks on the feature side of the wafer repeat steps 3-5 only washing the feature side
7. Place wafer on the hotplate
   - Cover with glass petri dish
   - Cover with aluminum foil
8. Repeat steps 2-7 for each wafer

**Anneal (hot plate)**

1. Program (temp, time, ramp):
   - 150 C, 10 min, max
   - 45 C, 1 sec, 120 C /h
2. Start the hotplate
3. When the hot plate has cooled bellow 40^o^C it can be removed from the hot plate and placed into a plastic petri dish

E) Silane Treatment

*Trichlorosilane is very toxic when inhaled. Work under dedicated hood! Read MSDS. This procedure will coat the master with a layer of silicone; so polymerized PDMS can be easier separated form the master.*

1. Place masters facing the middle in a desiccator
2. Put 40 uL of Trichlorosilane in a glass vial
3. Place the vial into the middle of the desiccator
4. Draw vacuum (do not breath in exhaust air, use hood!)
5. Leave in desiccator for 1 hr under vacuum under the hood

Fabrication Defects

**Features not sticking**

**Description –** After developing the photoresist some features may lift off or slide on the wafer surface. It may also look like a flaking of the SU-8 features in which case they will be pulled off when PDMS is removed from the wafer

**Causes** – There are a number of causes for this occurring. The most likely culprits are a dirty wafer or over-development.

**How to Solve –** To solve this, develop the photoresist for 8 minutes next time. This should be plenty of time (especially for features smaller than 30um tall). This will solve possible over-development issues.

Also use extra care in the cleaning step next time. This should leave the wafer with a cleaner surface. With the cleaner surface the resist will stick to the wafer properly.

**Thickness**

**Description –** When the feature heights are measured they are not the desired elevation.

**Causes** - The cause of this is either spinning to fast or to slow.

**How to Solve** – This problem is one of the easiest to solve. If the features are to thick spin at a higher speed the next time. If the features are to thin spin at a lower speed next time. The exact spin speed will have to be experimentally determined. Also remember not to change the first step on the spin coater as that step is not used to determine the final thickness but to make sure the entire wafer is covered with photoresist.

**Edge Sharpness**

**Description –** Features on the wafer after lithography are merged together instead of being distinct.

**Causes** – There are a number of causes to this. The most common are the features on the mask not being sharp enough and the mask was improperly placed onto the wafer during exposure.

**How to Solve –** To solve this, first check the mask. When looking at the mask features should appear sharp. There may be some blurring but features more than 10 um away from each other should be distinctly separate on the mask. If the features are not separate on the mask then contact printing company to order a new mask.

When exposing make sure the mask is in contact with the wafer. To make sure there is contact place the mask onto the wafer, turn on the vacuum, and then lightly pull up on the mask. If it stays put then you have contact.

Lastly, check what side of the emulsion mask has the features. Generally the Phillips lab orders masks so the features are down when you read the words at the top of the mask. To check, scratch each side of the mask near the outside of the black circle with razor blade. Whatever side the black is removed on is the side with the features. This side should be placed into contact with the wafer when exposing.

**White Reside on the Wafer**

**Description –** After developing and drying the wafer there are white streaks on the silicon surface.

**Cause –** This is caused by the photoresist not entirely washing off during development.

**How to Solve** – To fix this rewash the feature side of the wafer with developer and then ample IPA. Finally dry the wafer.

**Photoresist not Fully Covering Wafer**

**Description –** Photoresist does not cover the entire wafer after spinning.

**Cause** – Not enough photoresist placed onto the wafer prior to spinning or the wafer was not properly aligned onto the spin coater.

**How to Solve –** First make sure wafers are placed onto the middle of the chuck. A uniform spinning is important for proper photoresist coverage.

Also for the next masters pour more photoresist onto the silicon wafer. Make sure that the poured photoresist is in the center of the wafer. If excess photoresist is poured onto the wafer it will be spun off during the spinning.

**Bubbles and Streaks in the Photoresist**

**Description –** When looking at the wafer surface after spin coating the photoresist is not uniform.

**Cause** – The streaks and commits are caused by dust particles on the surface of the wafer. When the photoresist is spun with these particles still on the wafer surface it will result in a photoresist layer that is not uniform. Bubbles are caused during the pouring of the photoresist.

**How to Solve** – This goes back to wafer cleaning. The better the wafer is cleaned the less defects will be present after spin coating. Most of these defects are not important especially if they are not near features. Bubbles are important though because there is no photoresist of any thickness there. If the wafers photoresist is covered in bubbles clean this wafer by going through the single mask processing steps above while skipping exposure. This will result in a wafer with no features on it after development. Then the master fabrication process can be restarted.

To Prevent bubbles in the photoresist make sure the photoresist in the pouring container hasn’t been filled in the last half hour. After filling the containers are full of bubbles that will then be poured onto the wafer so be patient and wait for the bubbles to escape the photoresist.

**Misaligned**

**Description –** With a 2-mask device, features are not where they are desired.

**Cause** – This is caused by the second layer not being properly aligned onto the bottom layer thus features on the top layer are skewed to the side.

**How to Solve** – This problem is very difficult to solve. The only way to do it is to keep attempting the aligning process until features are finally properly aligned.

**2 Layer PDMS Device Fabrication**

*The two masters need to be clean. If especially the flow master is very dirty (from reusing), pour a thin layer of PDMS on it, let is polymerize, peal it off, master will be shiny like new.*

A) Mixing PDMS (Control Layer): Thick Layer

1. Blow air into a small plastic cup
2. Add 5:1 Elastomer Base then Elastomer Curing agent to the cup (Thick Layer)

|  | 1 | 2 | 3 | 4 | 5 |
| --- | --- | --- | --- | --- | --- |
| new master | 45 + 9 | 90 + 18 | 135 + 27 | 180 + 36 | 225 + 45 |
| reused master  (with PDMS ring) | 25 + 5 | 50 + 10 | 75 + 15 | 100 + 20 | 125 + 25 |

3. Mix the PDMS in the cup for, at least, 2 minutes
4. Turn off the mixer while the cup is still around it to avoid spraying the lab with elastomer (Mixer is out of the PDMS when turned off)

B) Mixing PDMS (Flow Layer): Thin Layer

1. Blow air into a small plastic cup
2. Add 20:1 Elastomer Base then Elastomer Curing agent to the cup (Spin Layer)
   - Add 10g of Elastomer Base for the first master and 5g per each additional master

| 1 | 2 | 3 | 4 | 5 |
| --- | --- | --- | --- | --- |
| 10 + 0.5 | 15 + 0.75 | 20 + 1 | 25 + 1.25 | 30 + 1.5 |

3. Mix the PDMS in the cup for, at least, 2 minutes
4. Turn off the mixer while the cup is still around it to avoid spraying the lab with elastomer (Mixer is out of PDMS when turned off)
5. Place a Kim Wipe over the cup
6. Peel off excess PDMS from the wafer

C) Vacuum Aspirate the PDMS

1. Place the cup containing PDMS into the PDMS desiccator
2. Turn on the pump and create a vacuum
3. As vacuum is reached the PDMS will bubble up in the cup
4. When the bubbles are near the top of the cup release the vacuum and the bubbles will recede
5. Allow the bubbles to rise, then lower them 2-4 more times
6. Remove the metal ring from bellow the desiccator
7. The PDMS should still bubble up but it should be slower with bigger bubbles now
8. When the PDMS bubbles near the top of the cup lift the desiccator and then hit it onto the table top (may need to hit it a few times to lower the bubble layer)
9. Repeat this hitting procedure (previous 2 steps) a few times until the bubbles no longer reach the top of the cup
10. Once the bubbles are no longer rising wait and watch the PDMS until bubbles no longer appear in the PDMS and only a liquid at the bottom of the cup is left (should take about 5 min)
11. Turn off the vacuum pump and remove the PDMS cup from the desiccator. Repeat this for the other cup containing PDMS

D) Pouring a Thick Layer

1. Pour PDMS (Thick Layer batch, Control Layer 5:1) into the desired mold, pour slowly, weigh to be precise (if new to the process)

| new master | 45 g |
| --- | --- |
| reused master  (with PDMS ring) | 25 g |

2. Place lid on the mold
3. Place the mold into the middle shelf of the box furnace set at 65C for 25 min (60C for 30 minutes)
   - Don’t stack containers
4. After the cook remove the mold from the furnace and let it cool, with lid still on

E) Spinning the PDMS Layer

1. Place paper/foil around the spinner for ease of cleaning
2. Program the spin coater (the following is optimized for 80um thick PDMS layer) (Look at PDMS spin speed chart in the back to determine for a different speed)
   - Step 1	30 sec, 1100 RPM, 100 RPM/sec
   - Step 2	0 sec, 0 RPM, 1000 RMP/sec
3. Dust the master surface with house air
4. Place master onto the center of the spin coater
5. Turn on vacuum
6. Pour Hershey kiss amount (~2.5mL) of PDMS (20:1 mixture) onto the center of the wafer
7. Cover the Spin Coater with black lid
8. Start Program by hitting green button
9. After the program has run turn off the vacuum and move the wafer back to its carrying petri dish
10. Allow the layer to reflow for 15 minutes by placing it on a level surface
11. Place the spun coated master into the box furnace set at 65C for 25 minutes (60C for 30 minutes)
12. After the cook, remove the mold from the furnace and let cool, with lid still on (the PDMS should be sticky but not deform when a finger is pressed onto the surface)
    - Another way to test the PDMS is to place a finger on the PDMS and then lift the finger. If the wafer sticks on the finger and then falls after a few seconds then it has been heated for the appropriate amount of time

F) Cutting out the Thick Layer

1. After the PDMS mold has cooled take a scalpel and gently cut around the pattern (cut a wide amount, avoid cutting around features or applying to much pressure and breaking the master)
2. Remove the PDMS mold form the master, take care not to rip the mold on removal
3. Trim the edges of the mold using the lift cutter
4. Cut out each device using the lift cutter
5. Place tape onto the feature side of the PDMS molds
   - DO NOT punch holes at this point

G) Aligning Layers (use stereomicroscope with light form top)

1. Remove the tape from the feature side of the Thick Layer PDMS devices
2. Under microscope, align the thick devices over the features on the spun coated layers
3. Place the mold into a box furnace set at 65C for 1.5 hours (Longer is ok)
4. After the cook remove the mold from the furnace and let cool, with lid still on

J) Cutting out sorter and punching holes

*Holes will be punched through both layers, the opening will only appear in the layer where a channel is perforated*

1. After the PDMS mold has cooled take scalpel and gently cut around the devices
2. Remove the PDMS mold gently from the master, take care not to rip the mold on removal
3. For large holes use for flow layer use 1.5mm punch
4. For small holes (1mm) use the 1 mm biopsy punch (Control Layer)
5. If the hole pieces remain in the mold after punching don’t tear them out as this could damage the mold around them
6. Instead use a solid puncher to remove the pieces from the mold (punch from the feature side but try to avoid touching anything else on the front)

K) Plasma Bonding

**Cleaning Prior to Bonding**

1. Tape is layered over both the front and the back of the PDMS casting
2. Glass slides and the glass holder in the plasma cleaner have water run over them
3. A drop of mild scrubbing soap is dropped on the glass pieces and is rubbed into both the front and the back of the glass slides
4. The soap is washed off using water
5. IPA is rinsed over both surfaces
6. Glass slides are air dried using pressurized air (can or house air) and immediately placed into the plasma cleaner
7. Remove tape from the PDMS mold and place it into the plasma cleaner with the side that is to be bonded (feature side) on the top

**Plasma**

1. Close the plasma cleaner and pull vacuum (100 to 300 mTorr)
2. Move the circle valve to the part open position (180 degrees from open) until there is a pressure of ~700 mTorr in the plasma cleaner
3. Turn on the plasma at the medium setting for the 30 seconds (look for a purple glow in the plasma chamber)
4. After 30 seconds immediately turn off the plasma and open the chamber in a fairly fast manor, but care full not to move the chips with the airflow

**Bonding**

1. Remove the glass holder from the plasma cleaner
2. Align the PDMS mold onto the glass slide (once the PDMS touches the glass it will be stuck there)
3. Gently press on the PDMS mold starting at a corner away from the features
4. Apply pressure outward from the original corner trying to ensure few if any bubbles are present between the PDMS and glass surface
5. Place the slide and PDMS into the furnace set at 65C for 1.5 hrs
6. After 1.5 hrs (can be longer) remove the device and let cool

Device Defects

**Dirty Slides**

**Description –** Dirt, streaks, or other contaminants on the glass slides used in bonding.

**Cause** – This is caused by improper cleaning procedures.

**How to Solve** – Wash again. Use more water and more IPA. If contaminants are still present after this use a different glass slide.

**Dirty PDMS**

**Description –** Debris in the PDMS. Clearly visible after the PDMS has cured.

**Cause** – Dirt getting into the liquid PDMS during device creation. This can happen at a number of different points.

**How to Solve** – Since this problem could have been caused during any number of the steps it is advisable to do each of the following. Prior to using the plastic cup clean it out by blowing with pressurized air. This will remove dust in the cup prior to pouring PDMS into it.

The blender may have cured PDMS on it. Removing this can also help reducing the amount of contaminants in the PDMS we are making.

Cover the cup containing the PDMS between processing steps. Between each of these steps, if the cup is open to air, dust will be landing in it. Covering it is the simple solution to that.

After device fabrication, use tape to clean both the top of the PDMS and the bottom of the glass. It is hard to tell where the dust particles are and chances are most will be in one of these two locations.

**Bubbles During Bonding**

**Description –** Bubble between the PDMS layer and the glass slide.

**Cause** – Pour bonding between the PDMS layer and the Glass slide.

**How to Solve** – Ensure both surfaces are clean. If the PDMS or the glass slide are dirty bonding problems can occur.

When pressing the 2 pieces together, start with your thumb on one corner of the device and then progressively apply pressure from that spot.

Try to complete the bonding process relatively fast. If enough time is spent the bond won’t be as strong or could fail. Due to this try to complete bonding within 1 minute.

**Bubbles in PDMS**

**Description –** Bubbles in the cured PDMS layer

**Cause** – Improper pouring or not enough degassing.

**How to Solve** – When pouring the PDMS into the mold pour in a corner away from features. Bubbles will generally form but by doing this they should be away from features. If they are close to features a pipet tip can be used to move them through the PDMS and away from features.

Also make sure that all bubbles are gone during the degassing step prior to releasing vacuum.

**No Bonding**

**Description –** Trying to bond PDMS multiple times.

**Cause** – Plasmaing PDMS 2 or more times.

**How to Solve** – Once PDMS has been through the plasma to attempt to plasma clean it must be bonded. It cannot be re-plasma-bonded. As such if the first attempt doesn’t work then the device needs to be thrown away.

**Channels Collapsing**

**Description –** The top of a channel collapsed and is now in contact with the glass slide.

**Cause** – Applying too much pressure when plasma bonding.

**How to Solve** – When pressing the PDMS and glass slides together during plasma bonding not much pressure is required. Too much pressure will cause the features to collapse.

**PDMS Sticking to Wafer**

**Description –** When attempting to remove the PDMS from the wafer the PDMS is sticking to the Silicon

**Cause** – Not enough silane used to destock the silicon.

**How to Solve** – The current wafer is no longer viable but for the next one make sure the wafer surface has gone through the silane procedure.

**Misaligned**

**Description –** The control layer is offset of where it should ideally be located.

**Cause** – Aligning of these two layer devices is done by hand and then inspected in the microscope.

**How to Solve** – If the 2 layers have been bonded together there is no solution. If that heat has yet to occur the top layer can be pulled off the bottom layer and realigned on the bottom one. Realigning this way can be done a number of times until dust gets in between the layers.

**Wrong PDMS Mixture**

**Description –** This problem is very hard to detect. It will result in layers deforming more or less then they should. This problem is easiest to see when cutting out thick layers. If the ratio is to high (15+ : 1) then the PDMS will stop cutting and instead will rip and tear as it is cut out of the master.

**Cause** – This is simply caused by the user failing to properly measure or mistaking the thin and thick layers when mixing.

**How to Solve** – Pay attention when mixing PDMS as it is one of the most important processing steps when creating these devices.

**Under Cooked Bonding**

**Description –** When looking at the control layer some of the channels are closed. This can also be seen when removing the wafers from the furnace prior to alignment. The top layer should be solid and the bottom layer should be sticky but not deform when a finger is pressed against it.

**Cause** – The layers are not cooked long enough prior to bonding.

**How to Solve** – There are a number of things that can be done to solve this. First always place partially curing devices on the second shelf in the box furnace. Second leave at least 1.5 inches between each petri dish. Third allow the layers to heat for longer, the exact time is very hard to determine and can only be done so through a lot of trial and error.

If bonding errors are still occurring try to leave the device out for 2 hours after alignment and then heat the device.

**Over Cooked Bonding**

**Description –** When removing the bottom layer out of the furnace and touching it the thin layer doesn’t deform or stick to the finger. This will also cause the valving to fail at much lower pressures.

**Cause –** This is caused by heating the layers prior to bonding for to long.

**How to Solve** – The only way to solve this is to heat the layers for a shorter amount of time.

If bonding errors are still occurring try to leave the device out for 2 hours after alignment and then heat the device.

**Setting up the sorter**

*Proper set-up of the sorter requires practice. The following procedure is optimized for the facilities in the Phillips Lab. When trained, the whole procedure takes about 20 min. A set-up sorter can be used from an hour to up to days at a time.*

*We use a stereo microscope (Leica M205 FA) equipped with a computer-controlled camera (Leica DFC 310 FX), although an inverted compound microscope with sufficient working distance should also work fine.*

*Lower end dissecting microscopes can also be used, depending on application, although special attention needs to be paid to the optics of the microscope and camera for the detection of low-expression fluorescent markers.*

1. Check device for Defects/Alignment
   - If Defects are present do not use the device
2. Tape above and below the devices and then remove the tape (cleaning step, optional)
3. Tape devices to microscope (optional, we do not usually do this)
4. Gather 6 valve tubes (the long ones on, 6 for Sorter), shot glass sized beaker with a bit of DI water and 3ml syringe (you can reuse that one)
   - Pull ~1 cm of water into the tube by attaching the syringe on the Lure valve and holding the end with the metal connector into the little beaker with water
   - Insert the metal tip into the proper hole in the PDMS device (following the order listed in the appendix), makes a “plop” noise (hardest part, see illustration at the end), do not push down further!!!
   - Disconnect the syringe, connect the valve tube to the solenoid tube (make sure the tube goes from the appropriate hole in the device to controlling valve, use schematic down below)
   - Tape tube to something steady (so the tubing isn’t pulled out)
   - Repeat this process for each of the valves
5. Turn on monitor + scope + Wago controller by using the switch on the power strip, the computer should already be on
6. For the WAGO:
   - There will be a series of lights flashing that will end will green lights
   - If there are red lights on after 20 seconds unplug and re-plug again
7. Check if the Blue Ethernet Wire is connecting the WAGO and the computer
8. Open Matlab
   - Open proper code
   - Run code
   - If the Matlab cannot connect to the WAGO, check if the Ethernet address is correct (see instructions on desktop background)
   - Follow instructions on the screen
9. When the code reaches the fill valves section
   - Open the pressurized-gas tank (the right, horizontal knob directly on the tank)
   - The pressure gauge directly next to the tank will show the pressure in the tank (200 psi is minimum for sorting)
   - The pressure gauge on the left shows the pressure going out of the regulator, set it to 20 psi with the green/white knob that says “increase” and “decrease”
   - Open the far left knob to let the air flow into the system
10. Allow the valves to fill with water
    - This may take a few minutes, you can follow the progress in the dark field mode, filled valves will “disappear”
11. When valves have filled increase the pressure in the valves to the desired run pressure (30 psi) and leave on for 30 seconds
    - This will stress the valves and if there are going to be valve failures they will generally occur at this point
12. Close the gas outlet on the tank regulator (Left most knob on the regulator)
13. Secure outlet tubing for each of the outlets
    - Place one outlet tube in each of the outlets
    - Take the end of each outlet tube and place it into the proper outlet container
      - Make sure the end of the tubing is below the device
14. Next Connect the Flush tubing (screw a lid with tubes going through it onto a falcon tube)
    - Place the flush container (the falcon tube) in a rack next to the scope on the left
    - Connect the flush liquid tube outlet to the device (the one with the metal connector)
    - Connect the flush air inlet tube (with the Lure connector) into the pressure inlet
      - This is the divided blue tube with 2 green caps on it, use one for the flush
15. Presort worm container setup
    - Worms should be in S-Basal, 5000 worms/ml
    - Screw lid with tubing going trough it onto worm falcon tube
    - Connect the presorted worm container liquid outlet into the device
    - Connect the presorted worm container air inlet into the pressure inlet
      - The other blue tube with green cap

Shake the tube carefully and place the tube in the same rack as the flush container, they should be on the same level to avoid effects of gravity

16. Continue the Matlab program
17. Turn the gas back on by opening up the gas tanks regulator
    - Turn flow/flush to 7PSI
18. The flush and worm flow will now reach the chip, remove air from the system by sorting and especially reflushing manually
19. Zoom the camera into the appropriate position/focus, move the chip to center the sorting channel
20. Turn GFP on for labeled worms
21. Turn lights off in the room
22. Manually control the worm movement
23. Or Switch to automatic sort if desired
24. After sorting, turn off the gas by closing the left knob on the regulator
25. Disconnect the flow tube form the valve control. press 5 and enter to start the automatic flushing (you might have to restart the code before you can do that)
26. All valves will open and worms will be flushed out the outlets, if already past the flush or go back up the flow tube, clearing out the chip
27. After all worms are cleared, lower the exit tube below the level of the chip, pull tubing out of the PDMS

Cleaning up:

1. Close the knob on the gas tank and wait for the air to leave the regulator
2. Watch both gauges to drop to 0 and then close the knob on the left to close the regulator.

**Tube Cleaning**

**In general:**

- Clean tubes directly after usage; prevents worms and goo from drying in
- Only clean the “wet tubes”, tubes for valving or air inlets (unless accidentally gotten wet) do not need to be cleaned regularly
- Do not remove the metal connectors that are used to insert the tubes into the PDMS chip
- Liquid waste from this procedure can be dumped in the sink
- Wear gloves and safety goggles

**Procedure:**

1. Fill the syringe labeled “bleach” with 0.6 % bleach
2. Fill the syringe labeled “H2O” with sterile water (autoclaved, purified water)
3. Connect bleach syringe with blunt syringe tip and syringe tip with tube
4. Hold the non connected tube end over waste container (some bleach will leak)
5. Push syringe to fill tube with bleach
6. Let sit for 1 min
7. Hold the other tube end above waste container, disconnect bleach syringe from tube
8. Use “air” syringe to push out the remaining bleach (3x), fill syringe with air while not connected to blunt syringe tip, the tip remains in the tube
9. Connect water syringe and push ca. 1.5 ml of water through the tube
10. Repeat step 8.
11. Repeat step 9.
12. Repeat step 8.
13. Use “air in a can” to remove remaining water, do not hold above waste container
14. Place in “clean tube” case

Appendix

how to insert valve tubing

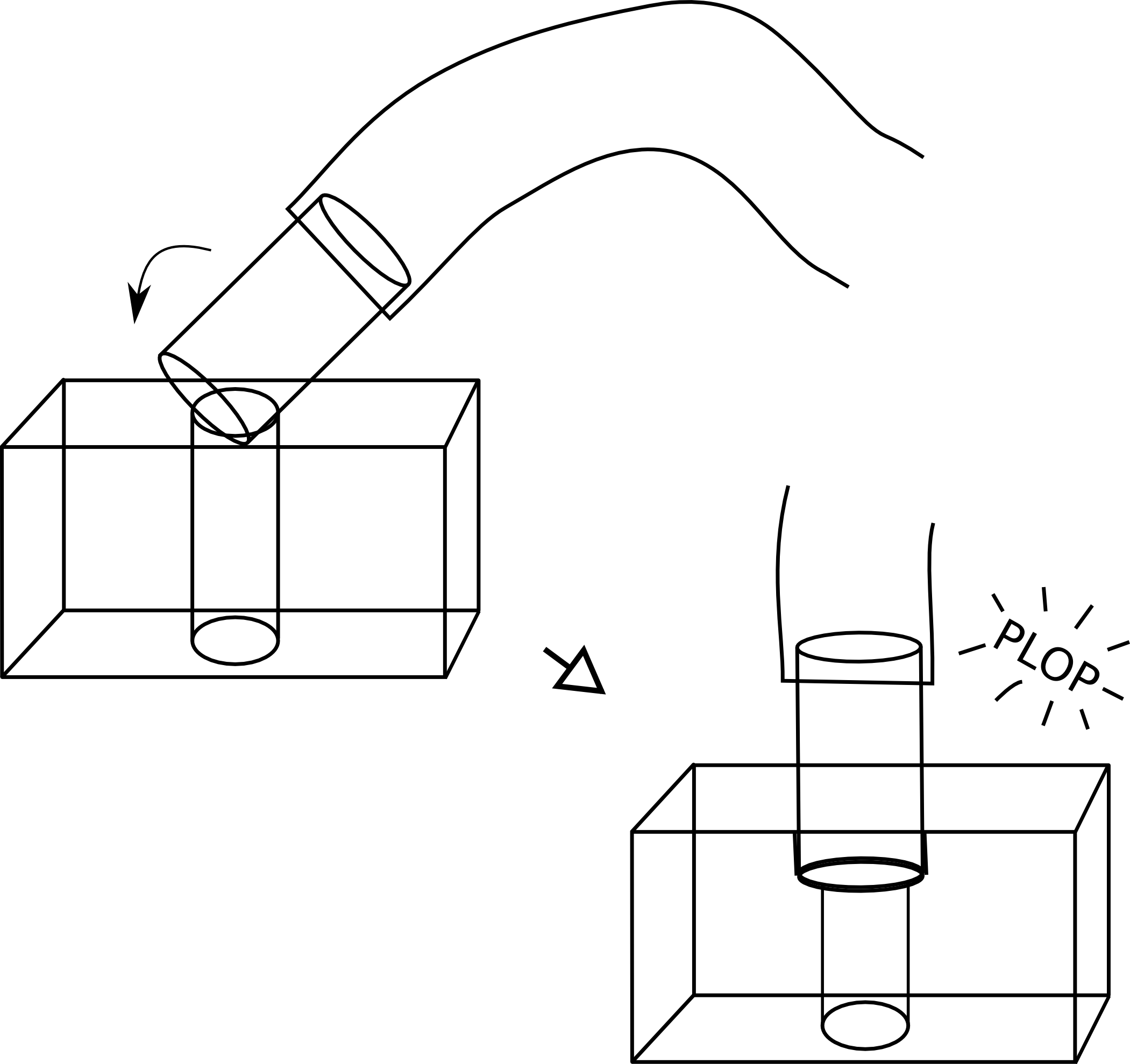

where to insert valve tubing

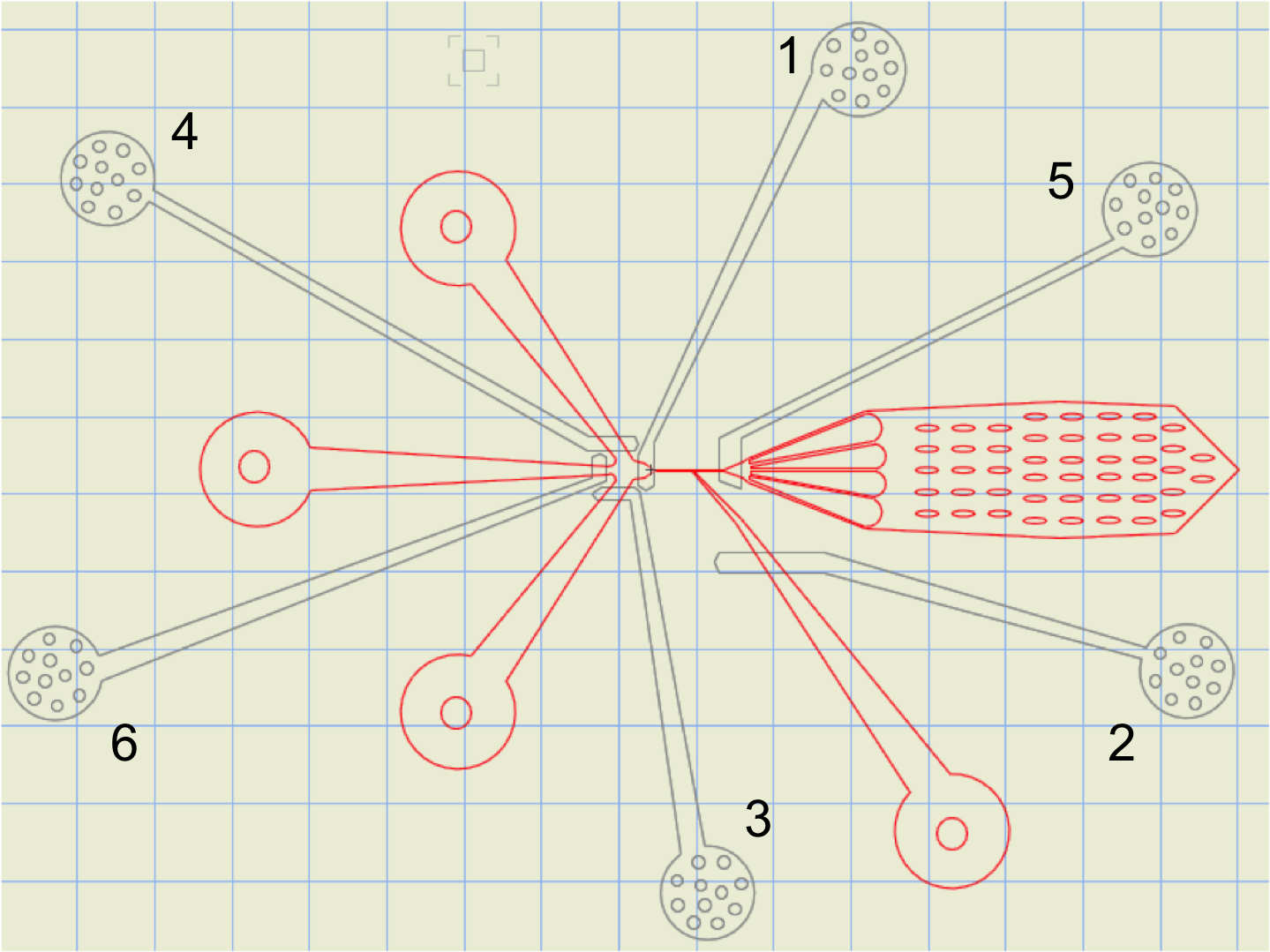
