## Supplementary material for "An open source microfluidic sorter for *Caenorhabditis* nematodes": S5 Parts List

Materials used for running a sorter and fabricating the devices

| Item | Company | Order size | order $ | used per set of 6 | per master set/chip |
| --- | --- | --- | --- | --- | --- |
| **Fabricating Masters** | | | | | |
| Wafer | University Wafers | 50 | 490 | 2 | 19.6 |
| SU8 2050 | Microchem | 500ml | 473 | 5ml | 5 |
| SU8 2050 | Microchem | 500ml | 504 | 5ml | 5 |
| High Petri Dish |  | 325 | 35 | 2 | 0.22 |
| CAD software | Vectorworks |  |  |  |  |
| Masks with design (reusable) | CAD/Art (local printing service) | 2 | 140 | 2 | 140 |
| SU8 developer | Microchem | 4l | 126 | 50ml | 1.6 |
| Labspace Photo-lithography | (local) | day use |  |  | 20 |
| Consumables |  |  |  |  |  |
| Time: for 2 sets of 6: 2h throughout the day | | | | | |
|  |  |  |  | Total: | 183.42 |
| **Fabricating Chips** | | | | | |
| PDMS | Sylgard | 500g | 67 | 7g | 1 |
| Glasslide | Fisher Scientific | 72 | 45 | 1 | 0.1 |
| Biopsypunches | Miltex | 25 | 114 | 0.3 | 1.4 |
| Scalpel | Miltex | 20 | 22 | 0.01 | 0.01 |
| Consumeables |  |  |  |  |  |
| Time: 2 sets of 6 chips: 2h throughout the day | | | | | |
|  |  |  |  | Total: | 2.51 |

| Item | Company | Amount | Prize each | Prize total |
| --- | --- | --- | --- | --- |
| **Sorter Hardware** | | | | |
| Pressure Regulator | McMaster Carr (6162k13) | 2 | 175 | 350 |
| Gauges | McMaster-Carr (4000K791) | 2 | 11.68 | 23.36 |
| Air Tank Regulator | VWR (55850-704) | 1 | 286 | 286 |
| Valve Controller | WAGO 1/0 end module (750-600) | 1 | 18 | 18 |
|  | WAGO Digital Output  Module, 4-channel  (750-504) | 1 | 59 | 59 |
|  | Wago controller (750-881) | 1 | 610 | 610 |
|  | Wago (750-1515) | 1 | 175 | 175 |
| Air Valve Array | Festo, Normally open: part# 197334, type MH1-A-24VDC-N-HC-10V-PR-K01-QM-AP-BP-CX-DX | 1 | 574 | 574 |
| Power supply | Digi-Key, CUI-ink  (VSK-520-24U-T) | 1 | 41.93 | 41.93 |
| DIN Rail | McMaster-Carr (8961K15) | 1 | 5 | 5 |
| tubing connectors and adapters | McMaster-Carr (5203K922) | 1 | 10.26 | 10.26 |
|  | McMaster-Carr (5779K131) | 1 | 8.12 | 8.12 |
|  | McMaster-Carr (7880T125) | 4 | 2.24 | 8.96 |
|  | McMaster-Carr (5779K44) | 1 | 4.75 | 4.75 |
| Ethernet cable |  | 1 |  |  |
| 3 prong plug |  | 1 |  |  |
| bread board |  | 1 |  |  |
| jumper cable |  | multiple |  |  |
| tubing | Pneumadyne PU-156F-0 | several meters |  |  |
| tubing | Scientific Commodities Inc. BB31695-P/9 | several meters |  |  |
| tubing | Pneumadyne PU-250PB-4 | several meters |  |  |
| Luer stubs adapter (17) | VWR (63019-820) | multiple | 244/ 100P |  |
| Luer adapters | McMaster-Carr (51525K32) | 6 | 4.82 | 28.92 |
| metal connectors | New England Small Tubes, 0.058” OD x 0.0475” ID x 0.500” Long | 9 |  |  |
| Fluorescence Dissecting Microscope with Camera | Scope: Leica M205FA & MDG41  Camera: Leica DFC310FX | 1 |  |  |
| Matlab software |  | 1 |  |  |
| image capture software | Micromanager | 1 |  |  |
| Computer to run software | DELL PRECISION T3610 WORKSTATION | 1 |  |  |
