## Supplementary material for "An open source microfluidic sorter for *Caenorhabditis* nematodes": S6 Code Scenario A: S6_ScenarioA_MarkedVsMarked.m

```
%%%%%%%%%%%%%%%%%%%%%%%%%%%%%%%%%%%%%%%%%%%%%%%%%%%%%%%%%%%%%%%%%%%%%%%%%%%
%%%  Sorting_Scenario A, Marked vs Marked                               %%%
%%%  Version 1.0                                                        %%%
%%%%%%%%%%%%%%%%%%%%%%%%%%%%%%%%%%%%%%%%%%%%%%%%%%%%%%%%%%%%%%%%%%%%%%%%%%%
%
% This program will be used to set up and sort nematodes manually or 
% automatically using a SORT 3rd generation devices. Worms will be sorted 
% left (Pharyngeal Marker), right(Coelomocyte Marker), or straight ahead.   
% To do this worms will be flushed into a sorting channel, imaged there, a 
% decision will be made, and then the worm is flushed into the correct 
% direction.
% 
%           Initial Setup Instructions
% 1) Power up the WAGO controller and plug it into the computer via the
% ethernet cable.
% 2) Test the connection between the controller and the computer by pinging
% the controller in the command prompt window of the computer. 
% If the connection fails check to make sure the wire
% connections are good and the IP address is correct. (update on line 37)
% 3) If there is still no signal check the controller and make sure there
% are no red lights on the control section of the WAGO
%
clc;
clear;
%%%%%%%%%% Variables %%%%%%%%%% 
male_counter=0;
female_counter=0;
waititeration=0;
holder = 0; % This value for user prompts
Worm_direction = 0; % direction the device will flush
con = 1; % user prompt to continue sorting
image_sort_counter = 0; % how many images are sorted before the user determines the next step
n = 0;  %Count

%%%%%%%%%% Connect with the WAGO and turn all valves on %%%%%%%%%%
clear('wago');
wObj = wagoNModbus('XXX.XXX.X.X', ones(1, 6), false); %Set correct IP Address

%%%%%%%%%% Valve Setup %%%%%%%%%%
holder = input('Do the valves need to be filled?\n1-Yes\n2-No\n');
if holder == 1
    fprintf('Please complete the following checklist.\nHit enter after completing tasks.\n');
    holder = input('Check for defects in the sorting device.  If defects are present please use a new sorting device.\n');
    holder = input('Fill control tubing with ~1cm of water.\n');
    holder = input('Place the control tubing into the correct inlets on the PDMS chips. Once all tubing is in place please continue.\n');
    wObj.setValves([0:5], ones(1, 6)); %This will turn all of the valves 6 on
    holder = input('wait until all of the valves are full of water.\n');
    wObj.setValves([0:5], zeros(1, 6)); %This will turn all of the valves 6 off
end
holder = input('Please connect Flush/Flow tubes.');
wObj.setValves(0, 1); % Standard valve setup conditions
wObj.setValves(1, 1);
wObj.setValves(2, 1);
wObj.setValves(3, 1);
wObj.setValves(4, 0);
wObj.setValves(5, 0);
wObj.setValves(6, 1);
wObj.setValves(7, 1);
wObj.setValves(8, 1);% End of Standard Valve Setup
fprintf('\nValves are set and manual testing is ready to begin. \n 1=Left, 2=Right, 3=Waste,  4=Begin Auto Sorting,  5=End Flush\n');

%%%%%%%%%% Manual Worm Sorting Setup-Testing Valves %%%%%%%%%%
while Worm_direction ~= 4
    
    Worm_direction = input('Direction -> ');
    
    %%%%% Worm Sorting Left %%%%%
    if Worm_direction == 1  
        wObj.setValves(4, 1); % Flush worms Left
        wObj.setValves(5, 1);
        wObj.setValves(2, 0);
        wObj.setValves(1, 0);
        wObj.setValves(0, 0);
        pause(0.25);
        wObj.setValves(0, 1);
        fprintf('Does the worm need to be reflushed? 1-Yes  2-No\n'); % Did worm flush?
        holder = input('->');
        while holder == 1 % Reflush Loop
            wObj.setValves(0, 0);
            pause(0.75);
            wObj.setValves(0, 1);
            fprintf('Again? 1-Yes  2-No\n');
            holder = input('->');
        end
        wObj.setValves(0, 1); % Return to Set position
        wObj.setValves(1, 1);
        wObj.setValves(2, 1);
        wObj.setValves(3, 1);
        wObj.setValves(4, 0);
        wObj.setValves(5, 0);
        
    %%%%% Worm Sorting Right %%%%%
    elseif Worm_direction == 2
        wObj.setValves(4, 1); % Flush worms Right
        wObj.setValves(5, 1);
        wObj.setValves(3, 0);
        wObj.setValves(1, 0);
        wObj.setValves(0, 0);
        pause(0.25);
        wObj.setValves(0, 1);
        fprintf('Does the worm need to be reflushed? 1-Yes  2-No\n'); % Did worm Flush?
        holder = input('->');
        while holder == 1 % Reflush Loop
            wObj.setValves(0, 0);
            pause(0.75);
            wObj.setValves(0,1);
            fprintf('Again? 1-Yes  2-No\n');
            holder = input('->');
        end
        wObj.setValves(0, 1); % Return to Set position
        wObj.setValves(1, 1);
        wObj.setValves(2, 1);
        wObj.setValves(3, 1);
        wObj.setValves(4, 0);
        wObj.setValves(5, 0);
        
    %%%%% Send Worm to Waste %%%%%
    elseif Worm_direction == 3
        wObj.setValves(4, 1); % Waste Valve settings
        wObj.setValves(0, 0);
        wObj.setValves(1, 0);
        pause(0.25); 
        wObj.setValves(1, 1);
        fprintf('Reflush the Sort Channel? 1-Yes  2-No\n') % Reflush Loop
        holder = input('->');
        while holder == 1
            wObj.setValves(1, 0);
            pause(0.75); 
            wObj.setValves(1, 1);
            fprintf('Again? 1-Yes  2-No\n');
            holder = input('->');
        end
        wObj.setValves(0, 1); % Return to set Postition
        wObj.setValves(1, 1);
        wObj.setValves(2, 1);
        wObj.setValves(3, 1);
        wObj.setValves(4, 0);
        wObj.setValves(5, 0);
        
    %%%%% End Manual Sorting %%%%%
    elseif Worm_direction == 4
        wObj.setValves(1, 1); % Close all valves but waste so more worms dont slip through
        wObj.setValves(2, 1);
        wObj.setValves(3, 1);
        wObj.setValves(0, 1);
        wObj.setValves(4, 1);
        
    elseif Worm_direction == 5
        wObj.setValves(0, 0);
        wObj.setValves(1, 0);
        wObj.setValves(2, 0);
        wObj.setValves(3, 0);
        wObj.setValves(4, 0);
        wObj.setValves(5, 0);
        pause(100); % 2 sec Flush
        
    else   % Catch invalid inputs
        disp('That is an invalid input.');
    end    
end

%%%%%%%%%% Auto Worm Sorting %%%%%%%%%%
wObj.setValves(0, 1); % Standard valve setup conditions
wObj.setValves(1, 1);
wObj.setValves(2, 1);
wObj.setValves(3, 1);
wObj.setValves(4, 0);
wObj.setValves(5, 0); % End of Standard Valve Setup


%%%%% Setting view area %%%%%
% some preliminaries
import java.awt.*;
rob=Robot;
conv=@(a,b,c) double(bitand(a,255^b))./255^c;
% define capture window size/pos (abs screen position)
xoff=0; % top left
yoff=0; % top right
xlen=1920; % width
ylen=1080; % height
% capture screen contents at this pos
rec=Rectangle(xoff,yoff,xlen,ylen);
cap=rob.createScreenCapture(rec);
dat=cap.getData;
buf=dat.getDataStorage;
% ...and create a ML var
buf=typecast(buf(:),'uint32');
buf=reshape(buf,dat.getWidth,dat.getHeight).';
mm=zeros([size(buf),3]);
mm(:,:,3)=conv(buf,1,0); % bitand(buf,255^1)./255^0;
mm(:,:,2)=conv(buf,2,1); % bitand(buf,255^2)./255^1;
mm(:,:,1)=conv(buf,3,2); % bitand(buf,255^3)./255^2;
mm=mm./255;
imwrite(mm, 'pica.jpg');
imshow('pica.jpg','border','tight');  % End of setting viewing area
[top_left_xp,top_left_yp] = ginput(1);
[bot_right_xp,bot_right_yp] = ginput(1);
top_left_y = round(top_left_yp); %input('top y? \n-');
top_left_x = round(top_left_xp); %input('left x? \n');
bot_right_y = round(bot_right_yp); %input('bot y? \n');
bot_right_x = round(bot_right_xp); %input('right x? \n');

xlen=bot_right_x-top_left_x;
ylen=bot_right_y-top_left_y;
xoff=top_left_x;
yoff=top_left_y;

holder = input('Viewing area is set.\nPlease close Figure 1.\n');
image_sort_counter = input('How many images are we sorting?\n');
fprintf('The computer will now take over sorting.\n');
tic;

    for counter = 1:image_sort_counter
        %%%%% Get Picture %%%%%
        pause(1)
         wObj.setValves(4, 1);
         import java.awt.*;
         rob=Robot;
         conv=@(a,b,c) double(bitand(a,255^b))./255^c;
         rec=Rectangle(xoff,yoff,xlen,ylen);
         cap=rob.createScreenCapture(rec);
         dat=cap.getData;
         buf=dat.getDataStorage;
         buf=typecast(buf(:),'uint32');
         buf=reshape(buf,dat.getWidth,dat.getHeight).';
         mm=zeros([size(buf),3]);
         mm(:,:,3)=conv(buf,1,0); % bitand(buf,255^1)./255^0;
         mm(:,:,2)=conv(buf,2,1); % bitand(buf,255^2)./255^1;
         mm(:,:,1)=conv(buf,3,2); % bitand(buf,255^3)./255^2;
         mm=mm./255;
         imwrite(mm, 'pica.jpg');
         
         
         
         
%%%%%%%%%%%%%%%%%%%%%%%%%%%%%%%%%%%%%%%%%%%%%%%%%%%%%%%%%%%%%%%%
% Fluorescent Marker Detection
% 
%%%%%%%%%%%%%%%%%%%%%%%%%%%%%%%%%%%%%%%%%%%%%%%%%%%%%%%%%%%%%%%%

% Reset all variables each loop (each time a picture is taken)
im=imread('pica.jpg');
maxValue=max(max(im));
      d = dir('pica.jpg');
         sorting_region_cols = [top_left_x, bot_right_x];
         region_width = sorting_region_cols(2)-sorting_region_cols(1); %Width of that regin
         too_many_region_limit = 6; 
            BW = im2bw(im,0.08);
            BW = imdilate(BW,strel('disk',25,0)); 
            rp = regionprops(BW, 'Area', 'Centroid'); %Compute Centroids
            n = length(rp); % Extract Centroids
            
             if (n <1)               
                    Worm_direction = 4;
                    
             else
            
                    centroids = reshape([rp.Centroid], [2,n]);
                    xs = repmat(centroids(1,:),[n,1]); % Compute the distance between all of the Centroids
                    ys = repmat(centroids(2,:),[n,1]);
                    dists = sqrt((xs-xs').^2 + (ys-ys').^2);
                    biggest_dist = max(dists(:)); % Largest distance
            
                   if (n > too_many_region_limit)
                      disp('Probably garbage too many Spots');
                      Worm_direction = 3;

                   elseif (n==1)
                             if (maxValue==255)
 
                                disp('myo-2');
                                Worm_direction = 1;
                             else
                                 disp('one dim dot');
                                 Worm_direction=3;
                             end

                   elseif (n>1)
                       if (maxValue<255)
                                disp('unc-122');
                                Worm_direction = 2;
                       else
                            disp('multiple dots but too bright');
                            Worm_direction = 3;
                            
                       end

                   else
                      disp('something weird'); 
                        Worm_direction = 3;
                  
                   end
             end
         
   
        
        %%%%% Wago Direction control %%%%%
        if Worm_direction == 1  %% Worm Sorting Left %%
            wObj.setValves(4, 1); % Flush worms Left
            wObj.setValves(5, 1);
            wObj.setValves(2, 0);
%             wObj.setValves(3, 1);
            wObj.setValves(1, 0);
            wObj.setValves(0, 0);
            pause(1);
            female_counter = female_counter + 1;
            wObj.setValves(0, 1); % Go to Set Position
            wObj.setValves(1, 1);
            wObj.setValves(2, 1);
            wObj.setValves(3, 1);
            wObj.setValves(4, 0);
            wObj.setValves(5, 0); % End of Valve Flush
          
        elseif Worm_direction == 2  %% Worm Sorting Right %%
            wObj.setValves(4, 1); % Flush worms Right
            wObj.setValves(5, 1);
            wObj.setValves(3, 0);
%             wObj.setValves(2, 1);
            wObj.setValves(1, 0);
            wObj.setValves(0, 0);        
            pause(1);
            male_counter = male_counter + 1;
            wObj.setValves(0, 1); % Go to Set Position
            wObj.setValves(1, 1);
            wObj.setValves(2, 1);
            wObj.setValves(3, 1);
            wObj.setValves(4, 0);
            wObj.setValves(5, 0); % End of Valve Flush
         
        elseif Worm_direction == 3  %% Send Worm to Waste %%            
                wObj.setValves(4, 1); % Waste Valve settings
                wObj.setValves(0, 0);
                wObj.setValves(1, 0);
                wObj.setValves(2, 1);
                wObj.setValves(3, 1);
                wObj.setValves(5, 0);
                pause(1); 

            wObj.setValves(0, 1); % Go to Set Position
            wObj.setValves(1, 1);
            wObj.setValves(2, 1);
            wObj.setValves(3, 1);
            wObj.setValves(4, 0);
            wObj.setValves(5, 0); % End of Valve Flush
                       
        elseif Worm_direction == 4 %% No Worm %%
              if waititeration < 3
                    wObj.setValves(4, 0);
                    waititeration=waititeration + 1;    
                    pause(0.3);
              
              else
                    waititeration=0;
                    wObj.setValves(4, 1); % Waste Valve settings
                    wObj.setValves(0, 0);
                    wObj.setValves(1, 0);
                    wObj.setValves(2, 1);
                    wObj.setValves(3, 1);
                    wObj.setValves(5, 0);
                    pause(0.5); 

                    wObj.setValves(0, 1); % Go to Set Position
                    wObj.setValves(1, 1);
                    wObj.setValves(2, 1);
                    wObj.setValves(3, 1);
                    wObj.setValves(4, 0);
                    wObj.setValves(5, 0); % End of Valve Flush

                end
           
              
        elseif Worm_direction == 6  %% Unplug the chip %%
            wObj.setValves(4, 1); % Waste Valve settings
            wObj.setValves(0, 0);
            wObj.setValves(1, 0);
            wObj.setValves(2, 1);
            wObj.setValves(3, 1);
            wObj.setValves(5, 0);
            pause(1);
            wObj.setValves(0, 1); % Go to Set Position
            wObj.setValves(1, 1);
            wObj.setValves(2, 1);
            wObj.setValves(3, 1);
            wObj.setValves(4, 0);
            wObj.setValves(5, 0);
            pause(0.2);
            wObj.setValves(4, 1); % Waste Valve settings
            wObj.setValves(0, 0);
            wObj.setValves(1, 0);
            wObj.setValves(2, 1);
            wObj.setValves(3, 1);
            wObj.setValves(5, 0);
            pause(0.5);
            wObj.setValves(0, 1); % Go to Set Position
            wObj.setValves(1, 1);
            wObj.setValves(2, 1);
            wObj.setValves(3, 1);
            wObj.setValves(4, 0);
            wObj.setValves(5, 0); % End of Unplug Flushes
        else 
            pause(0.5);
      
        end
    end
  
    fprintf('\n\n\n');
    disp('Pharyngeal Marked Animals');
    disp(female_counter);
    disp('Coelomocyte Marked Animals');
    disp(male_counter);
    toc;
    con = input('Sort Worms Again? 1-Yes 2-No ->');
    


%%%%%%%%%% Flush out device after use %%%%%%%%%%
fprintf('The sorter is being flushed out to ensure sorted worms are out of the device\n');
Worm_direction = input('Please prepare the device to be flushed out.  Hit 1 then enter when ready.\n');
Worm_direction = 1;
while Worm_direction == 1
    wObj.setValves(0, 0);
    wObj.setValves(1, 0);
    wObj.setValves(2, 0);
    wObj.setValves(3, 0);
    wObj.setValves(4, 0);
    wObj.setValves(5, 0);
    pause(100); % 2 sec Flush
    Worm_direction = input('Flush out device again? 1-Yes  2-No\n');
end
%%%%%%%%%% End Communication with WAGO and Camera %%%%%%%%%%
wObj.close();
clear wObj;
      
        
        

        
        
        
        
        
        
        
        
        
        
        
        
        
        
        
        
        
        
        
        
        
        
        
        
        
```
